# High Throughput Morphological Screening Identifies Chemically Defined Media for Mesenchymal Stromal Cells that Enhances Proliferation and Supports Maintenance of Immunomodulatory Function

**DOI:** 10.1101/2024.10.02.616329

**Authors:** Thomas M. Spoerer, Andrew M. Larey, Winifred Asigri, Kanupriya R. Daga, Ross A. Marklein

## Abstract

While mesenchymal stromal cell (MSC) therapies show promise for treating several indications due to their regenerative and immunomodulatory capacity, clinical translation has yet to be achieved due to a lack of robust, scalable manufacturing practices. Expansion using undefined fetal bovine serum (FBS) or human platelet lysate contributes to MSC functional heterogeneity and limits control of product quality. The need for tunable and consistent media has thus encouraged development of chemically defined media (CDM). However, CDM development strategies are rarely comprehensive nor considerate of a medium’s impact on MSC therapeutic function. Standard practice often neglects high-level interactions of media components, such as growth factors, that are critical to MSC growth and function. Given that MSC morphology has been shown to predict their immunomodulatory function, we employed a high throughput screening (HTS) approach to elucidate effects of growth factor compositions on MSC phenotype and proliferation in a custom CDM. This approach led to the discovery and refinement of several formulations that enhanced MSC proliferation and demonstrated wide ranging impacts on MSC immunomodulation. Overall, this work reflects how our novel HTS approach serves as a generalizable tool for the comprehensive improvement of MSC manufacturing processes.

## INTRODUCTION

Mesenchymal stromal cells (MSCs) have shown promise in preclinical studies as effective treatments for immune diseases^1–3^. These immunomodulatory effects are mediated by their secretome, which in part consists of a broad range of cytokines, chemokines, growth factors, and extracellular vesicles (EV)^4,5^. While using MSCs as an allogeneic therapy holds promise, achieving consistent and predictable quality during manufacturing is difficult due to a lack of standardized biomanufacturing practices^6–8^. Additionally, achieving the cell doses needed to support clinical trials (10^8^-10^9^ MSCs/patient) remains challenging, and significant scaling of biomanufacturing is necessary to obtain clinically relevant cell numbers^7,9^.

Traditional supplementation of MSC culture media with undefined components, such as fetal bovine serum (FBS) or human platelet lysate, can significantly contribute to MSC functional heterogeneity as the composition of these supplements varies considerably across batches^10,11^. Thus, there is great interest in development of chemically defined media (CDM) for MSC expansion to enable manufacturing of more consistent cell products. CDM can be defined as a growth medium wherein all components are identified and have their exact concentrations known. It has been demonstrated that the use of CDM reduces variability in MSC bioactivity across batches and consequently offers an avenue for standardization of MSC culture practices^12–14^. The use of CDM versus serum-containing media (SCM) has also been shown to enhance the proliferation of MSCs^13,15,16^. However, while several CDM formulations have been developed, the ability to understand the effect of the media and its individual components is difficult for academic researchers given that commercially available CDM formulations are proprietary. Secondly, the development of CDM is often laborious and incomprehensive as optimization of the myriad of variables that make up cell culture media is cost-prohibitive and challenging when using standard culture practices^17,18^. Culture media also constitutes a major part of the costs associated with MSC manufacturing, highlighting the need for development strategies that eliminate dispensable components and optimize for necessary ones.

The inclusion of growth factors is essential for supporting high cell proliferation rates in a manufacturing context^19,20^. Many growth factors including basic fibroblast growth factor (FGF), platelet-derived growth factor BB (PDGF), and epidermal growth factor (EGF) exhibit strong mitogenic activity on MSCs and impact their bioactivity^21,22^. Thus, several groups have screened the effect of different growth factors on MSC expansion. For example, Xu et al. screened the effect of 11 growth factors on MSC proliferation at predetermined concentrations by supplementing a CDM with each factor individually. After determining that FGF, EGF, and PDGF-AB significantly improved MSC proliferation in isolation, some combinations of these factors further increased cell growth in a synergistic manner^14^. Similarly, Jung et al. screened 4 different growth factors in several combinations to find an optimal formulation that enhanced MSC proliferation^23^. While these and other studies show unique growth factor combinations impart differential effects on MSC growth, they are limited to supplementing CDM with one or few growth factors at a time^13,16,24,25^. It is important to consider how the high-level interactions of these factors affect not only MSC growth, but their therapeutically-relevant functions i.e. immunomodulation. Therefore, there is a need to establish more comprehensive screening strategies for assessing the combinatorial effect of various growth factors on MSC growth and immunomodulatory function simultaneously for more robust CDM development.

Our group has previously demonstrated MSC morphology can serve as a strong predictor of immunomodulatory function (e.g. suppression of activated T cells) in response to priming with proinflammatory cytokines^26–28^. Measuring cell morphology is high-throughput and low cost, and it provides high-dimensional data at single cell resolution (‘morphological profile’) that can be indicative of complex intracellular states^29,30^. We previously used this tool in a high throughput screening (HTS) format to identify MSC priming conditions associated with unique morphological signatures and varied MSC-EV characteristics, demonstrating the value of leveraging morphology to guide manufacturing process modifications^31^. Thus, the goal of this study is to apply morphological profiling and high-content imaging (HCI) towards the comprehensive screening of growth factors supplemented into a chemically defined basal media (CDBM). The high throughput nature of morphological profiling afforded by HCI and automated image analysis enables screening of hundreds of different growth factor combinations and the identification of formulations that significantly affect MSC proliferation and phenotype. In this study, eight growth factors were screened in a full-factorial format, and media ‘hits’ were identified and applied towards MSC expansion over several passages to observe their effects on growth and immunomodulation.

## MATERIALS AND METHODS

### High throughput screening of MSC morphology

All MSC screening studies were conducted using either human adipose tissue-derived MSCs (Ad98), or human bone marrow-derived MSCs (BM71 and/or BM115) (RoosterBio). Donor information for all MSC lines used in this study can be found in **Supplementary Table S1**. MSCs were expanded to an initial population doubling level (PDLo, see **Table S1**), cryopreserved, and banked. Basal media formulations for CDM and SCM used in this study are outlined in **Supplementary Table S2**. Following a brief review of studies investigating CDM for MSC culture, eight growth factors were selected to investigate: fibroblast growth factor 2 (FGF), transforming growth factor β1 (TGF-β1), epidermal growth factor (EGF), insulin-like growth factor (IGF), platelet-derived growth factor BB (PDGF), leukemia inhibitory factor (LIF), stem cell factor (SCF), and Activin A (PeproTech). A two-level full factorial experimental design (2^8^ = 256 growth factor combinations) was used for screening (**Figure 1**). For exploratory and refinement screening, Ad98 (RoosterBio) was thawed, washed with αMEM basal media to remove residual serum components (Gibco), and seeded into 96-well plates (Corning) at a density of 1563 cells/cm^2^. Validation screening used all three MSC lines included in this study (Ad98, BM71, BM115). Wells receiving CDBM or CDM were coated with 2 μg/cm^2^ human fibronectin (Sigma-Aldrich) prior to cell seeding according to manufacturer’s instructions. For exploratory screening, Ad98 were either seeded in negative control conditions: 1) SCM or 2) CDBM, a positive control condition: SCM + 100ng/mL FGF, or a CDM treatment group. The design of experiments (DoE) tool in JMP Pro 16 was used to create the full factorial design and semi-random plate layouts.

**Figure 1:**
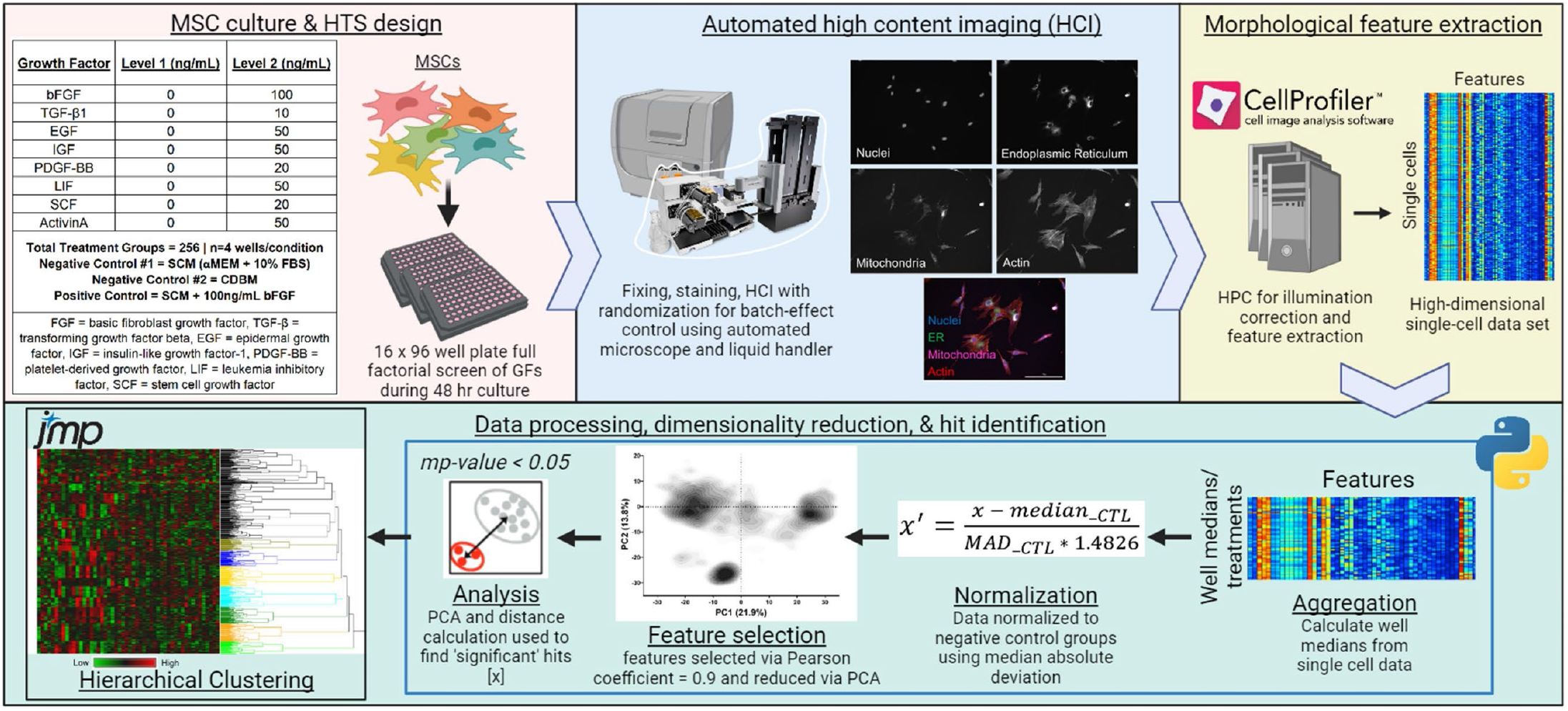
Schematic of HTS and morphological profiling approach. HTS of MSC morphological response to a combinatorial array of growth factors for CDM ‘hit’ identification using HCI and a modified Cell Painting protocol.

After 48 hours of culture, a modified Cell Painting^32^ protocol was performed (as previously described^31^) to assess MSC cellular and sub-cellular morphology via fluorescence microscopy and high content imaging (HCI). Briefly, MSCs were treated with MitoTracker Deep Red mitochondria stain (Invitrogen) for 30min at 37°C followed by fixing with 4% paraformaldehyde (PFA) (Electron Microscopy Sciences) for 20 minutes. After washing 2X with HBSS (Gibco), cells were stained with HBSS + 1% (w/v) bovine serum albumin (Sigma-Aldrich) + 0.1% (v/v) TritonX-100 (Sigma-Aldrich) + 8.25nM Phalloidin/AlexaFluor 568 conjugate F-actin cytoskeleton stain (Invitrogen) + 1.5μg/mL Wheat Germ Agglutinin Alexa Fluor 555 conjugate Golgi and plasma membrane stain (Invitrogen) + 5μg/mL Concanavalin A/Alexa Fluor 488 conjugate endoplasmic reticulum stain (Invitrogen) + 10μg/mL Hoechst nuclear stain (Invitrogen) for 30min, and cells were finally washed 3X with HBSS. All liquid handling for Cell Painting (e.g. fixing, staining, washing) was performed using a BioTek MultiFlo FX automated liquid handler (Agilent).

MSCs were subsequently imaged at 10X magnification using a centered 6×6 non-overlapping tile montage (% area of well imaged = 54%). HCI was conducted using a BioTek Cytation5 automated microscope (Agilent). Images were processed and morphological features were quantified using a combination of modified open source Cell Painting CellProfiler^33^ pipelines and Python scripts (Cell Painting GitHub, PyCytominer GitHub)^34,35^, as previously described^31^, using a high-performance computing cluster (HPCC) (Georgia Advanced Computing Resource Center). Briefly, images were first processed via an illumination correction pipeline (S1) and morphological features were quantified (S2) to generate high-dimensional single cell morphological data. A representative segmented image can be seen in **Figure S1**. Single cell output was then processed using modified PyCytominer Python scripts^35,36^ (S3, S4) to 1) aggregate single-cell profiles into well median profiles (S3), 2) normalize each well median profile feature to on-plate negative controls (SCM and/or CDBM) using median absolute deviation 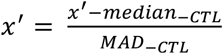 (S3), and 3) perform feature selection via Pearson’s correlation coefficient to remove features with a correlation coefficient >0.9 (S3). Principal component analysis (PCA) on correlations was then performed using all remaining features using JMP Pro 16.

Hit identification was performed leveraging cell count data and morphological profiles to discover growth factor combinations that significantly enhance MSC proliferation and influence MSC phenotype. Cell count data for each treatment were averaged and compared to the average cell count of each relative on-plate negative control (SCM and CDBM). In parallel, a Python script (Cytominer-eval GitHub)^36^ (S4) was used to collect principal components (PCs) that represent 90% of the variance in the data to conduct a multidimensional perturbation value (mp-value) analysis, as described previously^37^. This method assessed the significance of phenotypic differences between treatment groups and negative controls (SCM and CDBM) on a per-plate basis. To classify as a potential hit 1) cell count data for the treatment group must have a CV < 0.2 (to ensure consistency across replicates), 2) treatment groups must show significantly higher proliferation than both negative controls, and 3) morphological profiles of treatments must be significantly different from those of negative controls. PCA was run on this collection of potential hits, and PCs summarizing 90% of the variance in the data were collected, which was used for Ward’s hierarchical clustering of all potential hits. Final hits were identified by 1) ensuring all 4 replicates were in the same cluster (indicating low variability) and 2) choosing the treatment group with the highest proliferation per cluster with CV < 0.2. Representative images for visual quality control of hits were gathered using a Python script (S5) that identifies images with cell shape features closest to the average of well median replicates. This process was implemented to identify a suite of 8 unique hits that represent the wide landscape of morphological and desired proliferative responses to culture in screened CDM formulations.

### MSC expansion in control and media hits

Cryopreserved vials were thawed at 37°C, resuspended in αMEM, and centrifuged at 300g for 5 min. The cell pellet was washed with αMEM basal media, centrifuged, resuspended in respective hit or control media, and seeded in T-75 flasks (Greiner Bio-One) at a seeding density of 1,500 cells/cm^2^. Flasks receiving CDBM or CDM were coated with 2 μg/cm^2^ human fibronectin (Sigma-Aldrich) prior to cell seeding. Full media exchanges were performed using respective media types (prepared fresh) every 72 hours. Upon achieving 70%-80% confluence, MSCs were passaged into 3 replicate T-75 flasks at 1,500 cells/cm^2^, and remaining cells were resuspended in cryopreservation media (SCM + 10% DMSO) and cryopreserved for subsequent functional assays. For passaging, cells were washed 1X with phosphate buffered saline (PBS), harvested using 1X TrypLE (Gibco), washed with αMEM basal media, centrifuged at 300g for 5 min, and the cell pellet was resuspended in the respective expansion media type. MSC viability and cell counts were then acquired by AO/PI staining and automated cell counting (Nexcelom Cellometer K2).

### Indoleamine 2,3-dioxygenase (IDO) activity assay

Cryopreserved MSCs for each group were thawed, washed with αMEM basal media, resuspended in their respective expansion media type, and seeded into a T-75 flask at 4,080 cell/cm^2^. Cells were recovered for 48hrs, with the respective media type being fully exchanged after 24hrs. After 48hrs, MSCs were harvested using TrypLE and plated into 96 well plates at a seeding density of 40,000 cells/cm^2^ using either the experimental group’s respective expansion media type or SCM. After 24hrs of culture in 96 well plates, the medium was replaced in each well with protein-free RoosterCollect-EV media containing 50ng/mL IFNϒ and TNFα (termed ‘priming’). After an additional 24hrs, conditioned media (CM) was collected and stored at -20°C while cells were fixed with 4% PFA for 20 minutes, washed 1X with PBS, stained with 10μg/mL Hoechst nuclear stain (ThermoFisher) + 0.1% (v/v) Tween-20 (Sigma-Aldrich) for 1 hour, and washed 3X with PBS. Entire wells were imaged using a BioTek Cytation5 automated microscope, and Gen5 software (Agilent) was used for cell counting. CM samples were thawed and 100μL of each sample transferred to a 96 well plate (Corning). Samples were incubated with 30% Trichloroacetic acid (Sigma-Aldrich) to precipitate excess protein and centrifuged at 950g for 5 minutes. 75μL of supernatant was transferred to a separate 96 well plate and incubated with Erlich’s reagent for 5 minutes. L-kynurenine levels were determined based on absorbance (@490 nm) and normalized to cell numbers based on fixed nuclei count.

### Soluble intercellular adhesion molecule 1 (sICAM-1) ELISA

Additional CM samples (prepared identically to CM analyzed for IDO activity in previous section) were collected and stored at -80°C to assess levels of sICAM-1 using ELISA (Raybiotech). Briefly, all kit reagents and CM samples were thawed, and a standard curve was prepared. 100μL of standard solutions (triplicate) and CM samples (3 technical replicates) were added to pre-coated plates and incubated for 2.5hrs with gentle shaking at 70 rpm. Wells were washed 4X using a BioTek MultiFlo FX automated liquid handling machine, and 100μL of biotinylated antibody was added to each well and incubated for 1 hour with gentle shaking. Wells were washed 4X, and 100μL of Streptavidin solution was added to each well and incubated for 45 minutes with gentle shaking. Wells were washed 4X once more, and 100μL of TMB One-Step solution was added to each well and incubated for 30min in the dark with gentle shaking. Finally, 50μL of Stop Solution was added to each well and absorbance read at 450nm. sICAM-1 values were normalized to cell numbers based on nuclei count as described above.

### Statistical analysis

All statistical tests were conducted using GraphPad Prism 9. Specific statistical tests for each experiment are outlined in each figure legend.

## RESULTS

### Exploratory HTS identifies CDM formulations that produce distinct MSC phenotypes and enhance proliferation

**Fig. 1** presents the overall experimental design of the full factorial screen and tabulates the types and levels of growth factors studied, which included: FGF, TGF-β1, EGF, IGF, PDGF, LIF, SCF, and ActivinA (2^8^ = 256 total growth factor combinations studied). MSC phenotypic responses to media were compared to MSCs grown in SCM or CDBM as negative controls. The purpose of including CDBM was to directly observe how growth factor combinations may shift MSC growth and morphology, as direct comparisons to SCM may be confounded due to differences in the basal media formulations. SCM supplemented with 100 ng/mL FGF was included as an additional positive control on each plate. An array of statistical criteria was applied towards identification of growth factor combination hits. First, MSC proliferation in a CDM must be significantly higher than both the SCM and CDBM negative controls (**Fig. 2A**). This held true for 9 out of the 10 identified hits, where Hit J was chosen as a CDM that produced relatively similar proliferation and morphology to SCM. Second, treatment groups were considered hits if they were morphologically distinct from both negative controls based upon multidimensional perturbation value (*mp-*value) significance testing^37^, which returns a single test statistic based on multidimensional Mahalanobis distance between permutations of two groups using the number of principal components that represent 90% of the variance in the data (58 principal components) (**Fig. S2**). For Hit J, it was necessary for cell count and mp-value testing against SCM to yield a non-significant result. Hierarchical clustering (using 58 principal components) of hits that met the criteria was used to distinguish media that produced MSC morphologies unique from each other (**Fig. 2B**). Treatment groups with all four replicates in a cluster and the highest cell count of their cluster were chosen as final hits tabulated in **Fig. 2C**. Separate clustering was performed to ensure Hit J clustered with the SCM group (**Fig. S3**). Finally, Hit I was included to investigate whether over-supplementation of growth factors was unnecessary or even detrimental to MSC growth and function.

**Figure 2:**
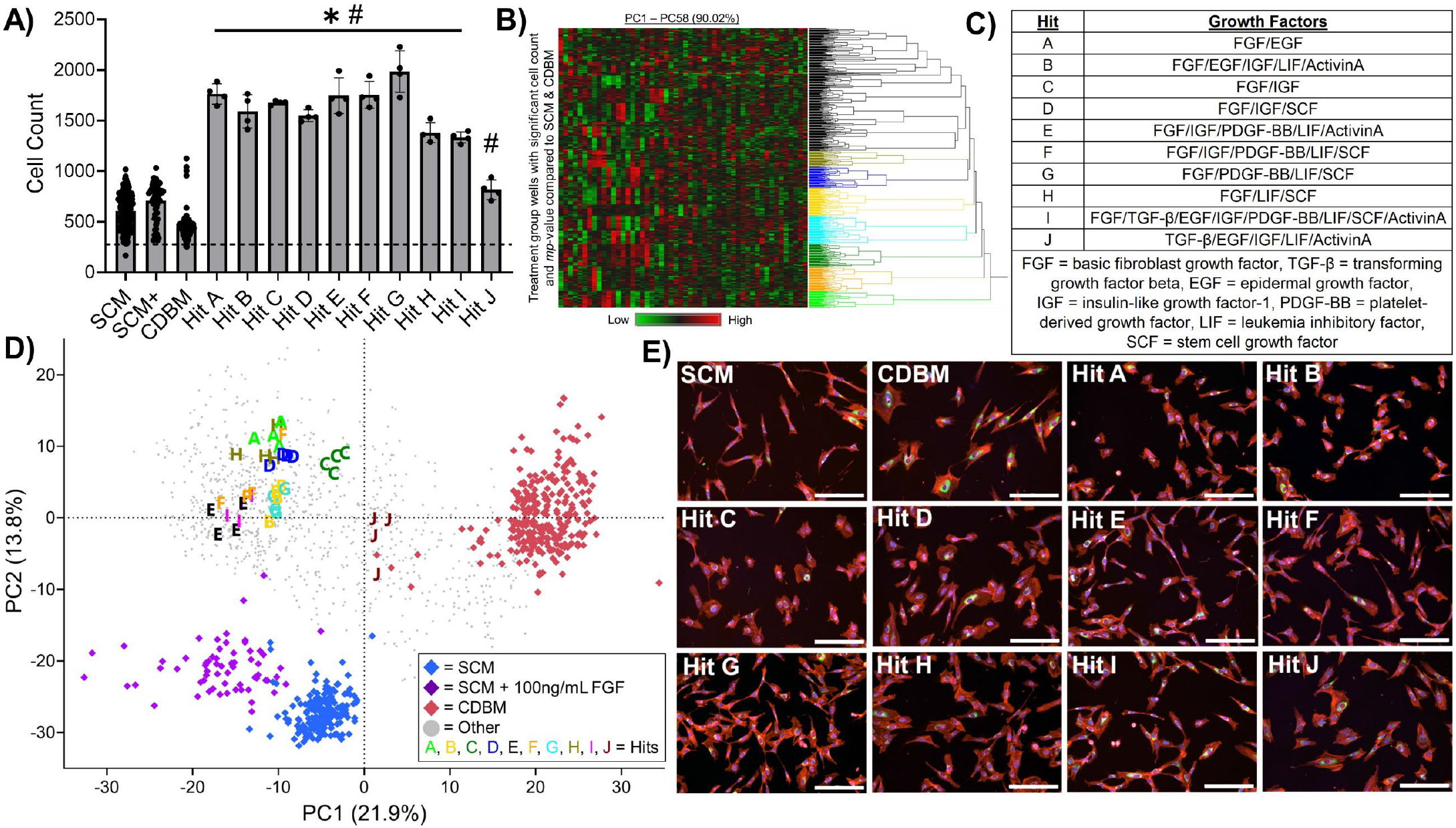
Exploratory morphological profiling HTS identifies growth factor combinations that induce distinct MSC phenotypes and enhance proliferation. A) MSC proliferation of hits and controls after 48 hours of culture where the dashed line represents initial seeding density. Cell count data from each media type was compared to SCM and CDBM on a per-plate basis via ordinary one-way ANOVA corrected for multiple comparisons via Dunnett’s test. \**p*<0.05 compared to SCM, #*p*<0.05 compared to CDBM. n=12 per plate for GM-MSC, n=12 per plate for CDBM, n=4 per plate for SCM+, n=4 for each treatment group. B) Hierarchical clustering of treatment groups that satisfy ‘hit’ criteria (i.e. significantly higher proliferation and distinct phenotype from controls based on significant mp-value (see Supplemental Figures)) distinguishes 8 hits from each other based on morphological principal components. C) Table of final identified hits to be included in subsequent validation screen. D) PCA plot representing morphological distinction between treatment groups and controls. E) Representative images from screen of MSCs grown in SCM and various hit media (red = actin, Golgi, plasma membrane, blue = nucleus, green = endoplasmic reticulum, magenta = mitochondria. Scale bar = 200μm).

Plotting the data using principal components 1 and 2 (35.7% of variance) further demonstrates uniqueness of hit media formulation morphologies as these groups cluster away from all controls and are well distributed along PC1 and PC2 throughout this high dimensional morphological space (**Fig. 2D**). As a means of assessing the quality of our HTS screen, plate heatmaps colored by PC1 and cell count were shown to not reveal patterns in terms of the hits and plate layouts (**Fig. S4**). This visual assessment suggests hits were identified based on replicable biological phenomena as opposed to experimental artifacts. Finally, visual inspection of representative images from each hit and control group served to qualitatively verify the effectiveness of this hit identification process to identify CDM formulations that produce distinct MSC morphological responses, as illustrated in **Fig. 2E**.

### Validation HTS confirms distinct morphological and proliferation response using multiple MSC lines

Hits A-J were included in a follow up screen to confirm the reproducibility of cell growth and morphological responses. The experimental design of this screen was identical to the exploratory screen; however, the SCM + 100 ng/mL FGF positive control was excluded, and two additional lines were included, BM71 and BM115, to further demonstrate the robustness of our screening approach and uniqueness of media hits. PCA plots of each cell-line show distinct morphological response to media hits from controls for all cell-lines (**Fig. 3A**). *mp*-value testing quantitatively verified that each hit is morphologically distinct from SCM and CDBM controls for all cell-lines (**Fig. S5**). However, PCA on combined morphological data demonstrates phenotypic distinction across donors, suggesting CDM hits do not decrease inherent heterogeneity of donors (**Fig. S6**).

**Figure 3:**
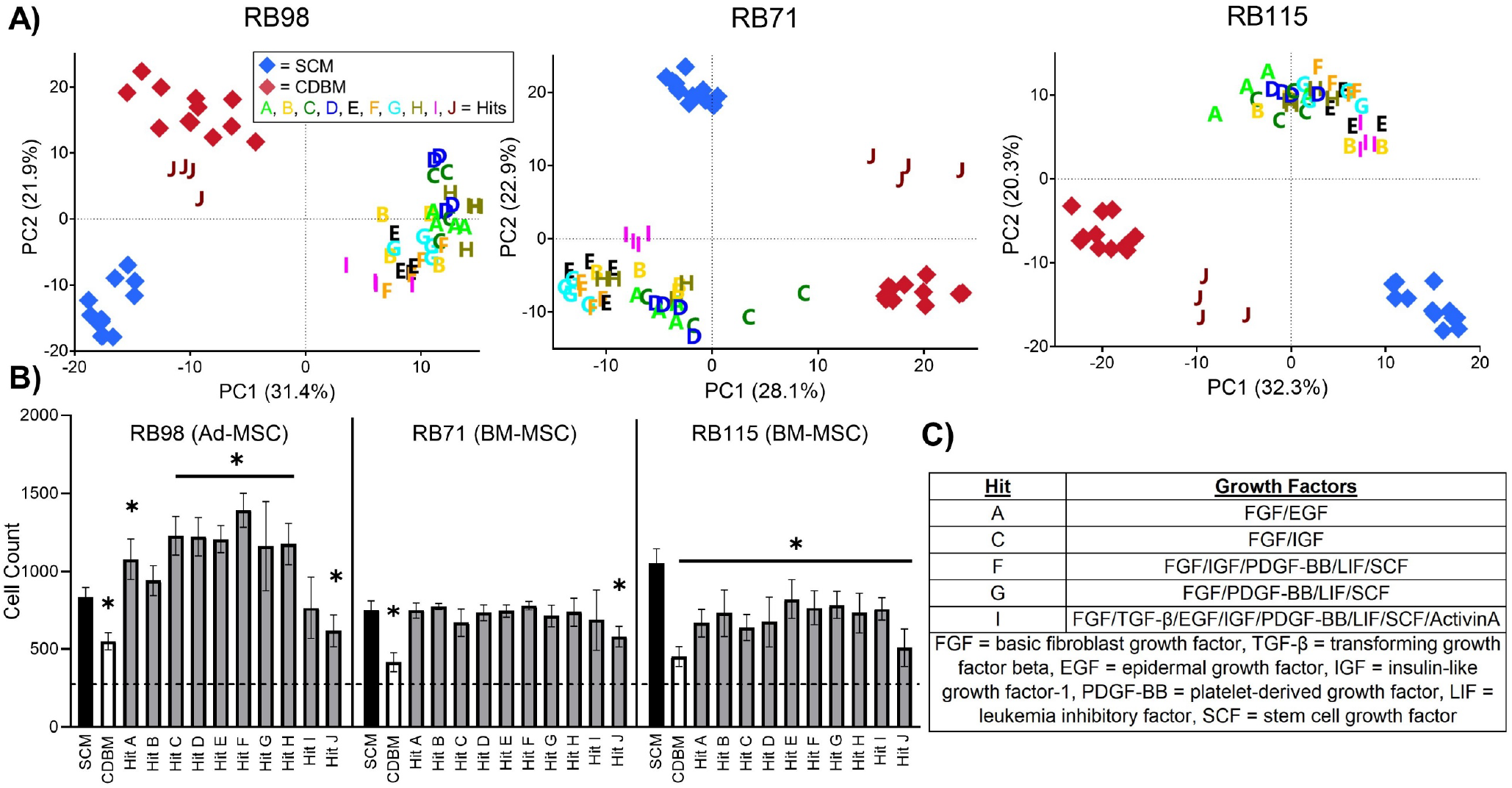
Validation HTS focuses scope to 5 growth factor combination hits, and different MSC donors show unique responses to same hit media. A) PCA plot for each cell line representing morphological distinction between hits and controls. B) Proliferation of hits versus controls after 48 hours of culture for each donor where the dashed line represents the initial seeding density for the screen. Cell count data from each media type was compared to SCM via ordinary two-way ANOVA corrected for multiple comparisons via Dunnett’s method. \**p*<0.05 compared to SCM of respective donor. n=12 per plate for GM-MSC, n=12 per plate for CDBM, n=4 for each treatment group. C) Table of final identified hits to be included in subsequent long-term expansion.

Proliferation data for Ad98 showed similar MSC growth across both the validation and exploratory screens, wherein most CDM hits show significantly higher proliferation than SCM (**Fig. 3B**). Notably, Hit B and Hit I did not show significant cell count differences in the validation screen (**Fig. 3B**) but did in the exploratory screen (**Fig. 2B**), which excluded Hit B from future experiments. Hit I was included again to investigate the effect of supplementing all growth factors on MSC expansion and function. Both BM71 and BM115 showed diminished proliferation compared to Ad98 when grown in hits (**Fig. 3B**). Together with morphological data, this finding motivated the use of Ad98 for subsequent experiments while also underscoring the need for further media refinement to enable identification of a universal CDM that produces consistent results across MSC lines. Multiple CDM formulations were considered hits based upon identical criteria to that used in the exploratory screen. We then narrowed the hits based on visual inspection of the PCA plot and a desire to choose hits that represent a diverse set of growth factors (**Fig. 3A & 3C**). Again, visual HTS quality assessment supported that hits were identified based on replicable biological phenomena (**Fig. S7**).

### Hits enhance MSC proliferation across multiple passages and maintain immunomodulatory function

Ad98 MSCs were cultured over three passages in SCM and each of the five CDM hits identified from exploratory and validation screening (**Fig. 4A**). Since the HTS was conducted over a period of only 48 hours in 96 well plates, we sought to investigate how media hits impact MSC yield and immunomodulatory function using a larger manufacturing scale (T-flasks) and duration (multiple passages/weeks). All CDM hits showed a higher fold change in cell yield compared to SCM after passage 1, which was consistent with findings from both the exploratory and validation screens (**Fig. 4B**). Notably, Hits F and G produced remarkably higher fold change than SCM and other CDM hits after 3 passages. However, it is evident that MSC proliferation in most CDM hits tended to decrease more dramatically over subsequent passages than SCM, especially after the first passage

**Figure 4:**
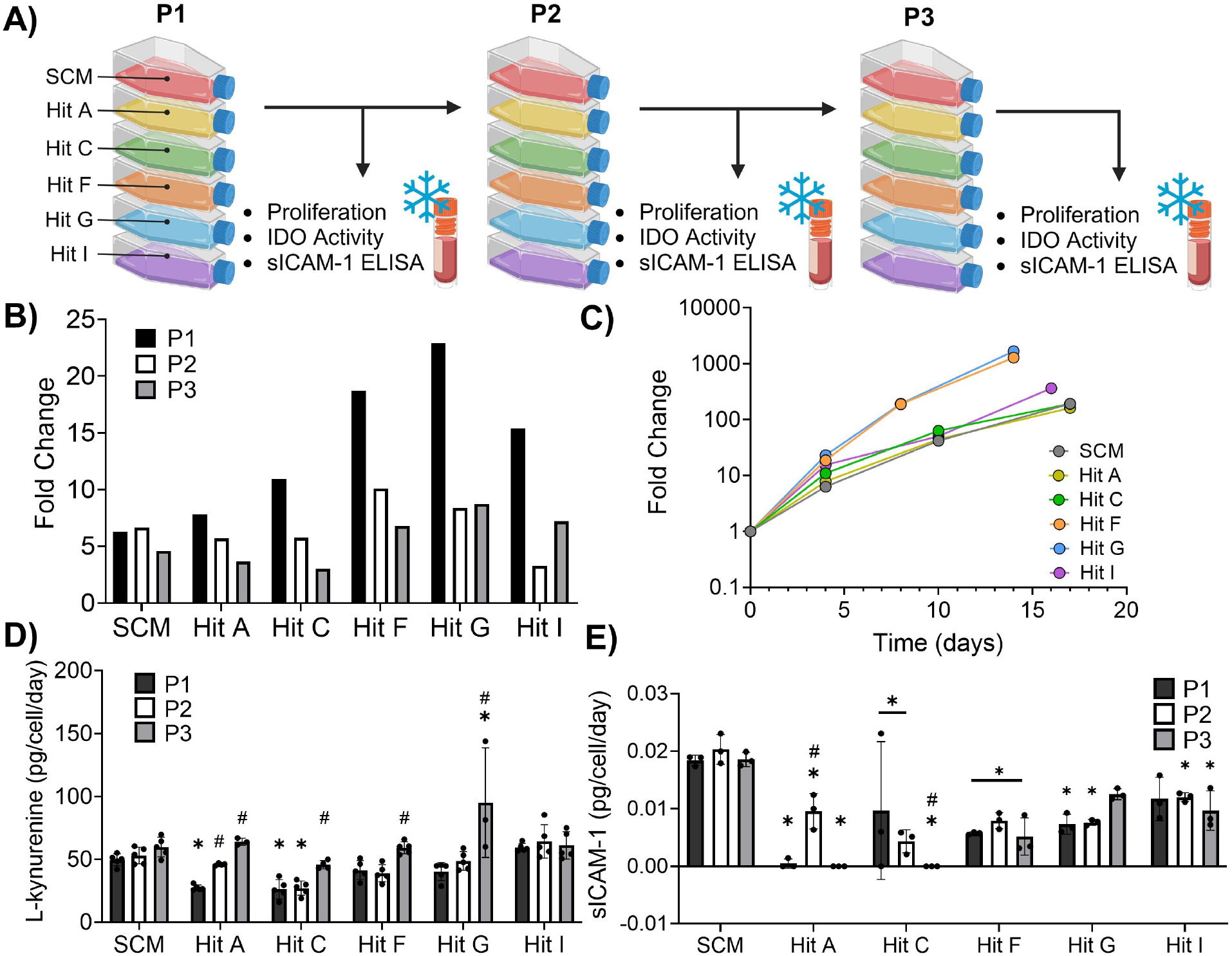
Select CDM hits enhance MSC proliferation and maintain immunomodulatory function across multiple passages. A) Schematic representing how expansion was conducted and assays performed at each passage. B) MSC proliferation fold change for each media type and passage. C) Cumulative fold change of MSC proliferation overtime for each media type. D) IDO activity measured by L-kynurenine levels normalized to cell number for each media type and passage; analyzed using ordinary two-way ANOVA with Dunnett’s post-hoc test (n=3-5 technical replicates, *p<0.05 different than SCM at P1, #p<0.05 different than P1 within media-type). E) sICAM-1 secretion normalized to cell number for each media type and passage; analyzed using ordinary two-way ANOVA with Dunnett’s post-hoc test (n=3 technical replicates, *p<0.05 different than SCM at P1, #p<0.05 different than P1 within media-type).

(**Fig. 4B**). Additionally, MSC viability at each passage was >80% for all media types (data not shown). It is important to note that in an industrial biomanufacturing setting most harvested cells would be passaged into additional flasks instead of immediately banked as was done here. Therefore, the cumulative fold change reflects the theoretical total MSC yield garnered from expansion in the different media types surveyed (**Fig. 4C**). Hits A and C showed similar cumulative fold change to SCM but have the added benefit of being chemically defined, while Hits F, G, and I stand out as producing markedly higher MSC yields in less time compared to SCM and other hits (**Fig. 4C**).

Most hit media did not significantly affect MSC immunomodulatory function across the 3 passages in terms of IDO activity, which has previously been correlated with MSC suppression of T-cell proliferation, especially upon priming with pro-inflammatory cytokines (**Fig. 4D**)^38,39^. Hits F, G, and I do not show any significant difference in L-kynurenine production per cell compared to SCM at each respective passage with the exception of P3 MSCs cultured in Hit G, which was the only CDM group to show significantly higher IDO activity than SCM (**Fig. 4D**). Of note, early passage MSCs grown in Hits A and C showed significantly less IDO activity compared to cells grown in SCM for the same passages (**Fig. 4D**). It was observed across most CDM groups that IDO activity generally increased with passage (Hits A, C, F, and G) whereas IDO activity remained consistent across passages for SCM and Hit I. Secretion of soluble ICAM-1 has similarly been shown to correlate with MSC’s ability to suppress cytotoxic T-cells and was therefore used as another indicator of MSC immunomodulatory function^26^. In this case, it was found that almost all CDM formulations significantly decreased MSC secretion of sICAM-1 compared to SCM at each passage (**Fig. 4E**), suggesting potential detrimental effects of hit media on MSC T-cell modulation. Of note, cells grown in Hit G at P3 and Hit I at P1 did not show any significant difference in sICAM-1 secretion compared to SCM of the same respective passage (**Fig. 4E**). Importantly, changes to growth factor concentrations in hits had yet to be considered, and the relatively high levels of each factor used for screening could contribute to the observed decrease in sICAM-1 secretion^40^. Therefore, one CDM hit formulation (Hit G) was chosen for further optimization due to the following considerations: 1) MSCs had the highest cumulative proliferation in Hit G, 2) Hit G was the only media to produce cells with significantly higher IDO activity than SCM, 3) MSCs grown in Hit G had sICAM-1 levels similar to SCM at the latest passage.

### Refinement HTS identifies lower growth factor combinations in optimal CDM formulation

Morphological profiling was used to screen the effects of altering the concentration of each growth factor included in the Hit G formulation. It was hypothesized that formulations which generate similar MSC proliferative and phenotypic responses to Hit G (but with lower growth concentrations) will produce similar MSC yields and potentially increase immunomodulatory function with added benefit of reducing cost. Thus, each growth factor in Hit G was tested at 2 levels in a full factorial format and compared to both SCM and the original Hit G formulation as a positive control (**Fig. 5A**). The experimental design of this screen was identical to the validation and exploratory screens. PCA of the high dimensional morphological data illustrates how cells grown in hit media possess distinct morphological signatures compared to the negative controls (**Fig. 5B**). In this case, almost all treatment groups cluster with the positive control, Hit G (+CTL) even for media formulations with the lowest concentrations for each growth factor (**Fig. 5B**). This is confirmed by *mp*-value testing, which shows no significant morphological differences between hits and +CTL but does show significance compared with negative controls (**Fig. S8**). Likewise, cells grown in refined hits show similar and even enhanced proliferation compared to +CTL (**Fig. 5D**). Cell counts observed for MSCs grown in +CTL are also similar to what was observed in both the validation and exploratory screens, which demonstrates reproducibility of our screening approach and consistency of CDM formulations. Although several treatment groups could be considered hits (no significant difference or significantly higher proliferation and non-significant *mp***-**value compared to +CTL), the final refined hits (Hit[Low] and Hit[Med]) were chosen based on the fact that both produced significantly higher proliferation to +CTL and significantly lowered the cost of the CDM (**Fig. 5C**). Hit[Low] was selected because it had the lowest concentration of each growth factor (and thus lowest cost). Hit[Med] showed the highest proliferation in the screen while also presenting similar morphology to +CTL. Again, visual HTS quality assessment and representative images supported that hits were identified based on replicable biological phenomena (**Fig. 5E, Fig. S8**).

**Figure 5:**
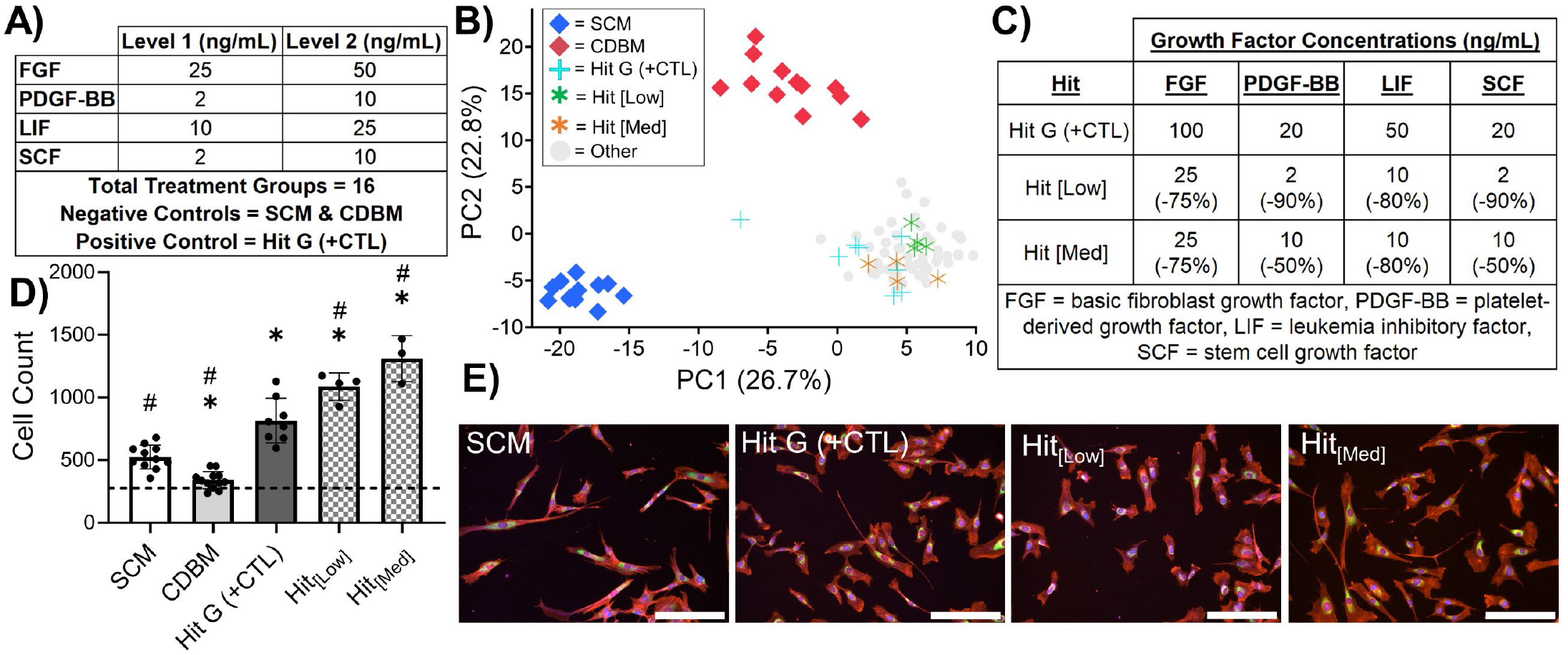
Morphological screening identifies CDM formulations with lower growth factor concentration without compromising phenotype or growth. A) Full factorial design parameters for screening growth factor concentrations. B) PCA plot representing morphological comparison between treatment groups and controls. C) Table of final identified hits to be included in subsequent expansion where the value in parenthesis reflects percent change in concentration for each growth factor. D) MSC proliferation of identified hits versus controls after 48 hours of culture where the dashed line represents the initial seeding density. Cell count data from each treatment group was compared to SCM on a per-plate basis via ordinary one-way ANOVA corrected for multiple comparisons via Dunnett’s test. \**p*<0.05 compared to SCM, #*p*<0.05 compared to Hit G (+CTL). n=12 per plate for SCM, n=12 per plate for CDBM, n=8 per plate for Hit G (+CTL). n=4 for each treatment group. E) Representative images from screen of MSCs grown in various media (red = actin, Golgi, plasma membrane, blue = nucleus, green = endoplasmic reticulum, magenta = mitochondria. Scale bar = 200μm).

### Optimized CDM hits maintain immunomodulatory function and enhance proliferation of MSCs

To confirm optimized hits with reduced growth factor concentrations produced proliferation and immunomodulatory function in MSCs similar to MSCs expanded in Hit G (with maximal growth factor concentrations), Ad98 MSCs were cultured in Hit G (+CTL), Hit[Low], Hit[Med], and SCM for one passage. MSCs expanded in +CTL and SCM exhibited similar fold change in proliferation compared to passage 1 of the previous expansion (**Fig. 4B**) with cells grown in the +CTL CDM still showing markedly higher yield compared to SCM (**Fig. 6A**). Both Hit[Low] and Hit[Med] were similar to +CTL, suggesting these optimized formulations do not compromise MSCs proliferative capacity (**Fig. 6A**). MSCs grown in +CTL, Hit[Low], and Hit[Med] all demonstrated similar IDO activity whereas cells expanded in SCM showed significantly less IDO activity compared to +CTL (**Fig. 6B**). sICAM-1 secretion was similar across all groups with no notable differences (**Fig. 6C**). This suggests that using lower growth factor concentrations in the optimized hits does not negatively impact MSC immunomodulation.

**Figure 6:**
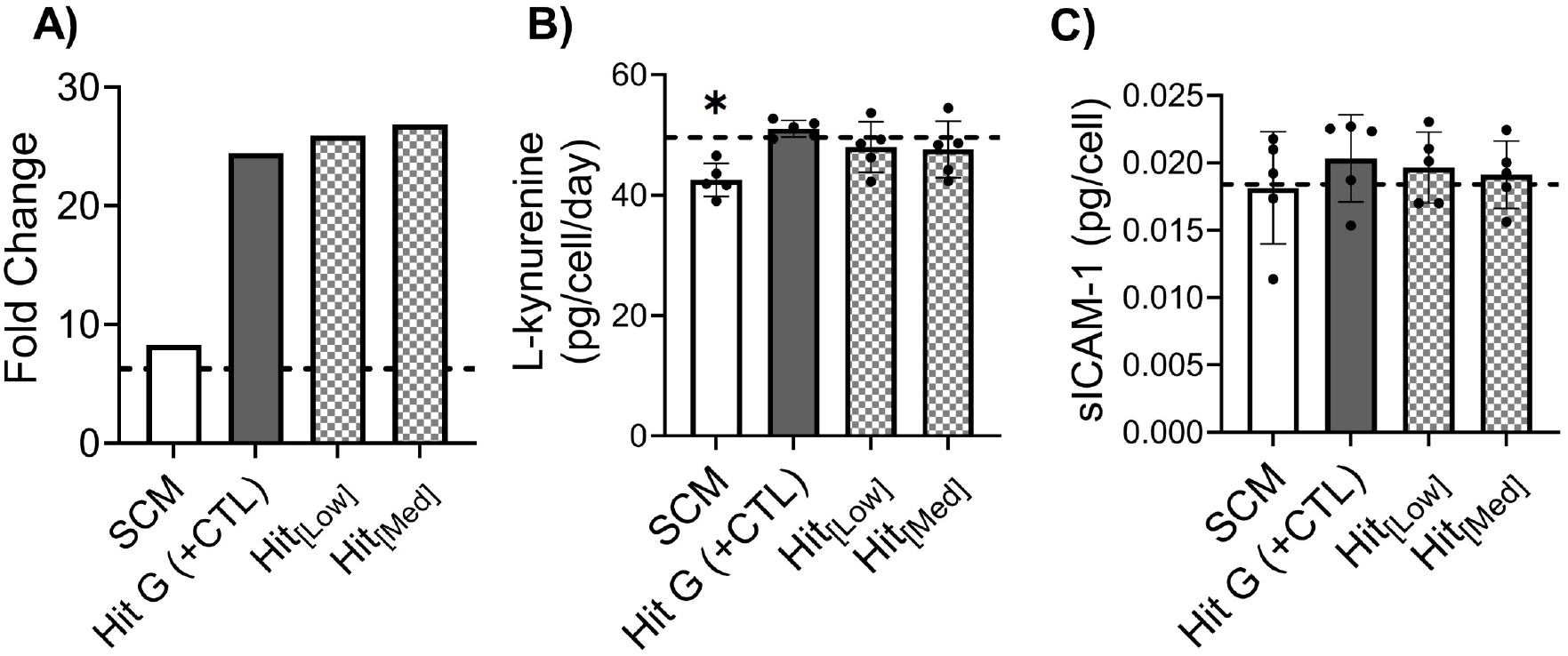
MSC expansion in optimized CDM validates maintenance of immunomodulatory bioactivity and enhanced proliferation compared to SCM. A) MSC proliferation fold change for each media type after one passage. B) IDO activity measured by L-kynurenine levels normalized to cell number for each media type; analyzed using one-way ANOVA with Dunnett’s post-hoc test to determine significance between hits and +CTL (n=5 technical replicates). C) sICAM-1 secretion normalized to cell number for each media type; analyzed using one-way ANOVA with Dunnett’s post-hoc test to determine significance between hits and +CTL (n=5 technical replicates). The dashed line represents respective values for SCM after P1 of the initial expansion (i.e. data presented in Figure 4). \**p*<0.05 compared to Hit G (+CTL).

It is important to note that cells grown in +CTL at P1 performed similarly to the SCM group in this expansion while they did not in the initial expansion (**Fig. 4E**). The significantly decreased sICAM-1 secretion observed in the initial expansion (**Fig. 4E**) prompted us to interrogate our method for conducting functional assays (IDO and sICAM-1 secretion) and investigate whether exposure to different media at certain points in the assay timeline affected functional outcomes. It was found that after initial culture rescue of cells in their respective media type (hit or control), the media used to seed cells into the assay plate affected MSC’s response to priming with pro-inflammatory cytokines IFNϒ and TNFα (**Fig. S9**). **Fig. S9** demonstrates that seeding all cells with SCM for the assay (after expansion in their respective experimental media) produces more similar MSC responses to priming than seeding cells in their experimental media. This suggests that short term exposure of MSCs to different media before priming may significantly impact their immunomodulatory function and therefore could be an additional manufacturing step to optimize in the future.

## DISCUSSION

Supplementation of MSC culture media with FBS or human platelet lysate is a major source of functional heterogeneity in MSC populations as compositions of these supplements vary across batches^10,11^. The HTS approach applied in this study allowed for the identification of CDM formulations most likely to produce desired outcomes during long-term expansion: high proliferation and maintained immunomodulatory function. To our knowledge, this study demonstrates the most comprehensive screen of growth factor-mediated impact on MSC growth and function to date since other studies have been limited to studying individual supplementation or only a few different growth factor combinations^14,16,41^. In addition, other screening studies predominantly use single-variable outcomes, such as cell proliferation alone, to determine the selection of growth factors for incorporation into a CDM^14,42^. Contrastingly, this study showcases the ability of morphological profiling using the Cell Painting protocol to distinguish experimental groups based upon multidimensional, functionally relevant criteria in a high-throughput format. Despite the fact that Cell Painting stains intracellular structures rather than specific biomarkers, observed alterations in morphology have been widely associated with functionality across diverse cell types, including neural cell differentiation potential in neural stem cells^43^, metastatic potential in cancer cells^44,45^, and immunomodulatory function of MSCs^26–28^. Additionally, our group has previously demonstrated success in applying Cell Painting and high throughput screening to optimize EV production in bioreactors^31^.

The CDM hit media in this study generally resulted in a MSC morphology with smaller spread area, decreased elongation, and increased roundness compared to our standard SCM formulation. These trends are consistent with several other studies that qualitatively compare MSC morphology in media supplemented with growth factors, including serum-free media^12,46,41,47^, chemically defined media^14,16,48^, and FGF-supplemented media^49,50,51^ to SCM. However, cell-line dependent morphological differences persisted, wherein the validation screen showed CDM altered MSC morphology compared to respective controls for each cell-line, but the three MSC donors presented different morphological and proliferative responses to the same CDM. This suggests that either HTS using morphological profiling needs to be conducted using multiple cell-lines or CDM may best be developed on a specific cell-line/product basis.

Along with these observed changes in morphology, growth factor supplementation in CDM significantly enhanced MSC proliferation during screening and expansions, corroborating the results of many previous studies^13,14,16,21,22,25,41,52,53^. Relevant to this work, FGF and PDGF have been previously shown to augment MSC proliferation during early passages^49,50,54,55^, which is consistent with results shown in this study for the optimized Hit G media. The growth factors included in this study are known to be potent mitogens, binding to their respective transmembrane receptors and activating downstream pathways; in particular, the phosphoinositide-3 kinase (PI3K)-Akt/protein kinase B (PKB) and the mitogen-activated protein kinase (MAPK) Erk pathways. Related to morphology, relationships between focal adhesions, actin cytoskeleton rearrangement, activation of Rho kinase, and subsequent induction of growth factor-mediated cell cycle progression have been well documented^60–62^. Sustained activation of the ERK pathway by growth factor stimulation for G1 phase cell cycle progression is dependent upon actin cytoskeleton organization and actin-myosin contractility, which manifests in cell spreading in fibroblasts^61^. While MSCs grown in SCM showed larger cell area and more prominent actin stress fibers in this study, a higher proportion of smaller, more rounded cells found in CDM may be attributed to observation of more cells progressing through the mitotic G2 phase, wherein adhesion complex area decreases, and stress fibers disassemble^62^. Therefore, the consistent relationship between small, rounded morphology and enhanced cell cycle progression observed in this study and others may be expected, since cytoskeletal condensation and contraction are necessary cellular operations for mitotic division, triggered by aforementioned signaling pathways^63,64^.

The exploratory and validation screens showed that our HTS approach successfully identified hits with morphological signatures significantly different from the SCM and CDBM controls. Morphological signatures of MSCs grown in hits were also distinct from one another, which coincided with distinct cellular behavior in terms of growth and immunomodulation during long-term expansion. It is important to note that while morphological differences between MSCs grown in most CDM hits and SCM were apparent during screening and coincided with varied immunomodulatory function at each passage, cells grown in Hit G and the optimized CDM hits had similar immunomodulatory function to those grown in SCM. Thus, despite the fact MSCs in these refined media showed highly distinct morphology and proliferation, their IDO activity and ICAM-1 secretion did not vary significantly on a per-cell basis. However, when considering overall differences in cell behavior, or ‘quality’ (i.e. proliferative capacity, IDO activity, sICAM-1 secretion), between MSCs in refined hits and SCM, morphology did serve as an indicator of differential quality of cell populations isolated from these different media. Expansion in CDM generated higher yields of similarly bioactive MSC products to those expanded in SCM, which, in a clinical setting, would allow for administration of higher doses of similarly bioactive cells, thus increasing the overall effectiveness of administered product. Therefore, the overall manufacturing efficiency and product quality harvested from CDM deviated from that generated using SCM, which was captured by differential morphology during screening.

Importantly, expansion of MSCs across multiple passages showed that some CDM hits significantly elevated MSC proliferation while maintaining their immunomodulatory function. More specifically, Hits F, G, and I greatly boosted cumulative fold change of Ad98 compared to the SCM control, while other CDM hits showed similar growth to SCM but have the added value of being chemically defined. This result is expected since each of the selected hit media showed significantly greater proliferation during the exploratory screen compared to SCM. In addition, the inclusion of growth factors specific to Hits F and G, such as FGF, PDGF, LIF, and SCF, have been included in media formulations from other studies, likewise demonstrating improved MSC proliferation^47,48,65^. However, the specific combination of these four growth factors is unique, to the best of our knowledge, and outperforms other growth factor combinations based on our findings, even those that contain more factors, such as Hit I. This suggests that over-supplementation of a culture media does not necessarily improve or maintain MSC proliferation or immunomodulation, a finding observed by other groups^14,52,66^. In fact, combinations of certain factors have been observed to inhibit growth, such as combining ActivinA and TGF-β1 when culturing induced pluripotent stem cells (iPSC)^67^. Together, these findings further motivate the development of more specific, cost-effective CDM formulations.

The screening method used in this study effectively identified growth factors that interact synergistically with each other. Notably, all CDM formulations that showed comparable IDO activity to SCM (Hits F, G, I) contained growth factors included in the final optimized hit formulations: FGF, PDGF, LIF, and SCF. Prior studies have consistently observed a synergistic effect of growth factors such as FGF, IGF, PDGF, and TGF-β on MSC proliferation, which agrees with results from the exploratory screen described here, where formulations with these growth factors in combination outperformed those in isolation^14,46,68^. However, the comprehensive nature of the screening method used here allowed for the discovery of new synergistic interactions, such as those between LIF, SCF, FGF, and PDGF. In fact, a previous study developing CDM for primary bovine satellite cells conducted an effect summary to find that cell proliferation was most significantly influenced by interaction effects between LIF, IGF1, PDGF, and VEGF^66^. Upon testing these factors in isolation and in different combinations, it was found that the culture media necessitated the inclusion of PDGF and LIF to reach similar growth levels to FBS-supplemented media^66^. While each of these growth factors are known to activate MAPK/ERK and PI3K/Akt pathways, some factors in our optimized media formulation activate distinct pathways, such as JAK/STAT. For example, LIF and SCF can affect STAT3 signaling^69,70^, which can crosstalk with PI3K/Akt and MAPK/ERK signaling to promote activation of these pathways^71^. JAK/STAT activation by IFN-γ also mediates increased IDO expression by MSCs^38,72^; thus, similar activation of this pathway by LIF, SCF, and IFNϒ highlights the importance of including these factors in a media formulation to sustain MSC immunomodulation.

It was observed during the first expansion that most MSCs grown in CDM had reduced sICAM-1 secretion compared to SCM. Upregulated ICAM-1 (CD54) expression by MSCs exposed to pro-inflammatory cytokines is known as an integral component of their immunomodulation of T cells, generation of T regulatory cells, and expression of amyloid-β-degrading enzyme neprilysin in microglia^73–75^. Notably, it has been shown that ICAM-1 can act as a therapeutic paracrine factor in soluble form, and MSC-derived microvesicles (MVs) are enriched in ICAM-1 upon exposure to TNFα and IFNϒ^76,77^. Our results align with previous results where MSCs cultured in serum-free conditions show lower CD54 surface marker expression compared to those in serum-containing media^78^. Due to the undefined nature of FBS^79^ it is possible that lot-specific factors could enhance MSC response to IFN-γ and TNF-α. Despite the higher sICAM-1 produced by MSCs grown in SCM, inconsistencies and unknowns regarding FBS-mediated MSC functional response attest to the benefits of adopting CDM. It was also thought that the high growth factor concentrations leveraged for exploratory screening could inhibit ICAM-1 secretion, potentially via growth factor receptor desensitization or feedback inhibition. More specifically, suppressor of cytokine signaling protein (SOCS) upregulation by growth factor receptor binding has been postulated to participate in the negative feedback regulation of cytokine signaling pathways (e.g. JAK/STAT, NF-κB) important for ICAM-1 expression^80^. Upon initially observing decreased ICAM-1 secretion after expansion in CDM, we employed methodology to uncouple the biological effects of expansion media from priming media, which was used for the sICAM-1 assay, to observe how assay media impacts MSC response to priming. Results showed that the assay media likely plays a role in MSC response to priming but that the media used for initial expansion does not impact this response when assay media is kept constant across groups (**Fig. S9**). However, future morphological screening of assay media components could help clarify how CDM formulations impact MSC function when used as priming media, as this was deemed outside the scope of this study.

As a final demonstration of the utility of our HTS approach, we identified CDM formulations consisting of the same combination as Hit G but lower growth factor concentrations to reduce cost and potentially improve MSC growth and function. This screen successfully identified formulations with low growth factor concentrations that produced similar phenotypes to the positive Hit G control. The screen demonstrated MSC morphological and proliferative response to Hit G is consistently distinct from SCM and CDBM controls and is robust, as many formulations at lower concentration clustered with the positive control both in terms of growth and phenotype. Results after a one passage expansion in flasks aligned closely with those from the first passage of the initial expansion, where Hit G showed 24X fold change in growth, again highlighting consistent MSC responses to CDM across different manufacturing runs. Also, growth and immunomodulatory capacity for Hit[Low] and Hit[Med] were not significantly different from the Hit G +CTL, which again supports the notion that morphology after short-term culture can predict functional outcomes during manufacturing.

Overall, this work applies concepts traditionally used in drug screening to develop CDM for improved MSC manufacturing. This work represents the most comprehensive screen to date of growth factor-mediated effects in a CDM on MSC expansion and phenotype, which successfully identified growth factor combinations that increase proliferation while maintaining therapeutically relevant functions. This screening approach is generalizable towards other aspects of cell manufacturing processes/materials and can be readily adapted to different therapeutic cell-types. Optimization of other CDM components, including hormones, lipids, vitamins, etc., using the screening methodology described here would further refine the CDM formulations developed. Additionally, integration of more functionally relevant *in vitro* and *in vivo* assays would contribute towards understanding how CDM impacts disease-specific MSC functions. More studies are also warranted to uncover specific mechanisms and crosstalk between relevant pathways differentially activated by growth factors included in CDM. Finally, applying the CDM formulations discovered here to larger scale manufacturing platforms, such as bioreactors, is important for validating the consistency and translatability of this media for production of improved MSC products.

## Abbreviations

MSC: mesenchymal stromal cell
FBS: fetal bovine serum
CDM: chemically defined media
CDBM: chemically defined basal media
SCM: serum-containing media
FGF: basic fibroblast growth factor / fibroblast growth factor 2
PDGF: platelet-derived growth factor BB
EGF: epidermal growth factor
HCI: high content imagining

## Acknowledgements

Drs. Ty Maughon and Seth Andrews helped with the expansion and banking of all MSC lines used in the study. We consulted with Drs. Anne Carpenter, Beth Cimini, Shantanu Singh (Broad Institute) in the planning and development of our HTS approach. This study was supported in part by resources and technical expertise from the Georgia Advanced Computing Resource Center, a partnership between the University of Georgia’s Office of the Vice President for Research and Office of the Vice President for Information Technology. We consulted with Akash Ramachandran, Ravi Jyani, and Kailin Chen in modifying our HTS Python scripts and HPC execution.

