## Supplemental Figures and Tables for "High Throughput Morphological Screening Identifies Chemically Defined Media for Mesenchymal Stromal Cells that Enhances Proliferation and Supports Maintenance of Immunomodulatory Function"

| Cell-line | Tissue Source | Donor Age | Donor Sex | PDL |
|---|---|---|---|---|
| Ad98 | Adipose | 18-30 | Female | 12 |
| BM71 | Bone marrow | 18-30 | Female | 13 |
| BM115 | Bone marrow | 20 | Female | 15 |

**Table S1: Donor information.**

**A)**

| CHEMICALLY DEFINED BASAL MEDIA (CDBM) | | | | |
|---|---|---|---|---|
| **Component** | **Company** | **Catalog No.** | **Concentration (ug/mL)** | **Volume Ratio** |
| AdvancedMEM Basal Media* | Gibco | 12492013 | - | 97.78% |
| GlutaMAX | Gibco | 35050061 | - | 1.000% |
| Pen/Strep | Gibco | 15140122 | - | 1.000% |
| Chemically Defined Lipid Concentrate** | Sigma-Aldrich | L0288 | - | 0.100% |
| Putrescine | Sigma-Aldrich | P5780 | 10 | 0.100% |
| Hydrocortisone | Sigma-Aldrich | H0888 | 0.05 | 0.008% |
| Progesterone | Sigma-Aldrich | P8783 | 0.005 | 0.010% |

*https://www.thermofisher.com/order/catalog/product/12492013

**https://www.sigmaaldrich.com/US/en/product/sigma/l0288?srsltid=AfmBOopYkr4EzlWhiHCnkrpHshQfISV9e6hCNzv3olzRAzLelmTapDKh

**B)**

| SERUM CONTAINING MEDIA (SCM) | | | |
|---|---|---|---|
| **Component** | **Company** | **Catalog No.** | **Volume Ratio** |
| αMEM Basal Media | Gibco | 12492013 | 88% |
| GlutaMAX | Gibco | 35050061 | 1% |
| Pen/Strep | Gibco | 15140122 | 1% |
| Fetal Bovine Serum (FBS) | Neuromics | 218H19 | 10% |

**Table S2:** Media formulations for A) chemically defined basal media (CDBM) and B) serum-containing media (SCM). CDM hit formulations are CDBM supplemented with growth factors as defined in main figures. Full formulations for AdvancedMEM Basal Media and Chemically Defined Lipid Concentrate can be found via the links provided.

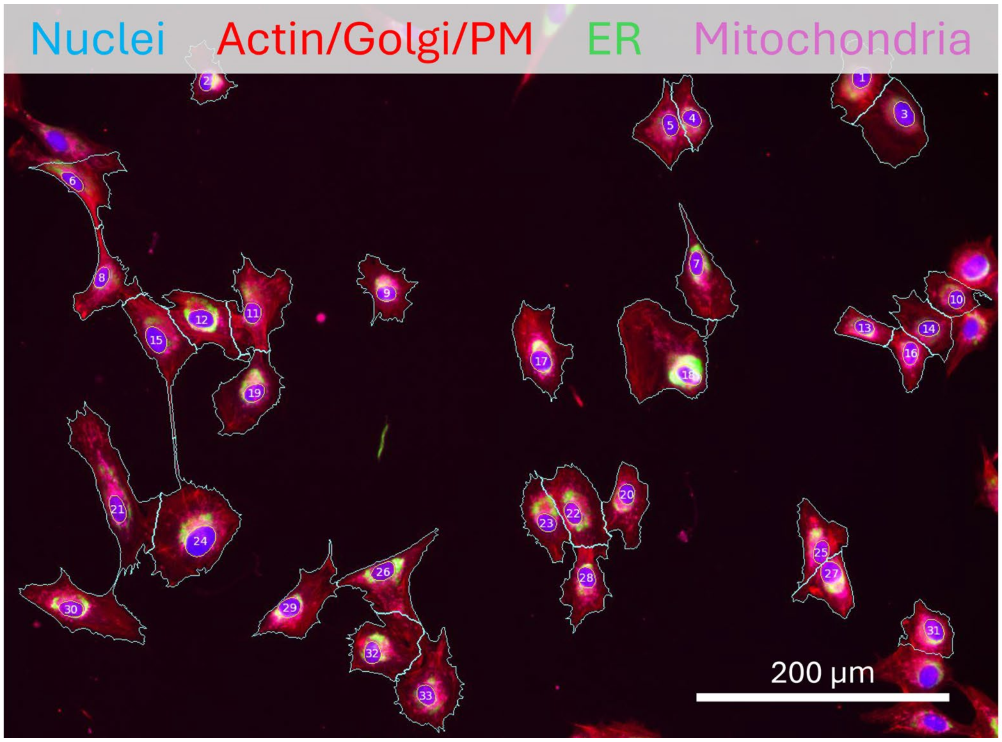


**Figure S1: CellProfiler pipeline accurately segments cellular and subcellular components for morphological profiling.** Representative color composite image of nuclear (yellow lines) and cellular (cyan lines) segmentation, with number in nucleus representing object count in CellProfiler metadata. PM = plasma membrane, ER = endoplasmic reticulum.

A)

| Treatment | Plate # | CDBM *mp* -value | SCM *mp* -value |
|---|---|---|---|
| CDBM | 3 | 0.999 | 0.004 |
| SCM | 3 | 0 | 1 |
| Hit J | 3 | 0 | 0.2 |
| CDBM | 4 | 0.999 | 0 |
| Hit F | 4 | 0 | 0 |
| SCM | 4 | 0 | 0.998 |
| CDBM | 5 | 0.999 | 0 |
| Hit H | 5 | 0 | 0 |
| SCM | 5 | 0 | 1 |
| CDBM | 6 | 0.999 | 0 |
| Hit D | 6 | 0.001 | 0.001 |
| SCM | 6 | 0 | 1 |
| CDBM | 9 | 0.997 | 0 |
| Hit B | 9 | 0 | 0.001 |
| SCM | 9 | 0 | 1 |
| CDBM | 10 | 1 | 0 |
| Hit G | 10 | 0.001 | 0.001 |
| SCM | 10 | 0 | 1 |
| CDBM | 12 | 1 | 0 |
| Hit I | 12 | 0 | 0.001 |
| SCM | 12 | 0.745 | 1 |
| CDBM | 13 | 0.998 | 0 |
| Hit E | 13 | 0.001 | 0 |
| SCM | 13 | 0.09 | 0.997 |
| CDBM | 14 | 0.999 | 0 |
| Hit C | 14 | 0 | 0 |
| SCM | 14 | 0.234 | 0.998 |
| CDBM | 15 | 0.994 | 0.006 |
| Hit A | 15 | 0.001 | 0 |
| SCM | 15 | 0.068 | 1 |

B)

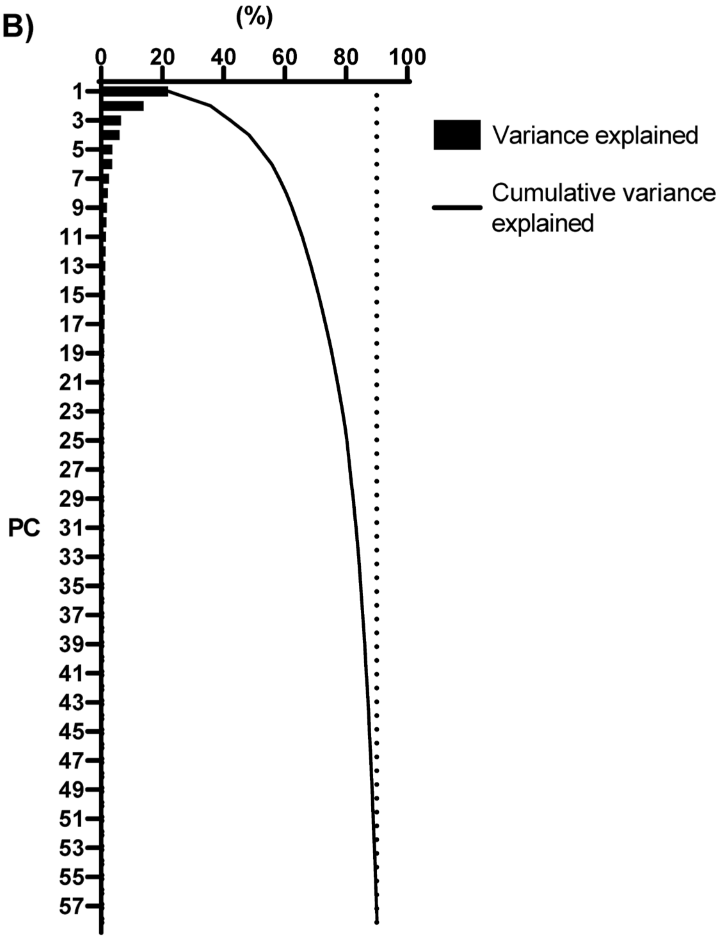


**Figure S2: Exploratory HTS mp-value testing highlights phenotypic differences between hits and controls using 58 principal components that represent 90% of the variance in the data.** A) mp-values for phenotypic statistical comparisons between each hit and on-plate controls. B) Exploratory HTS PCA cumulative variance plot showing number of PC's that represent 90% of the variance in the data.

**A)**

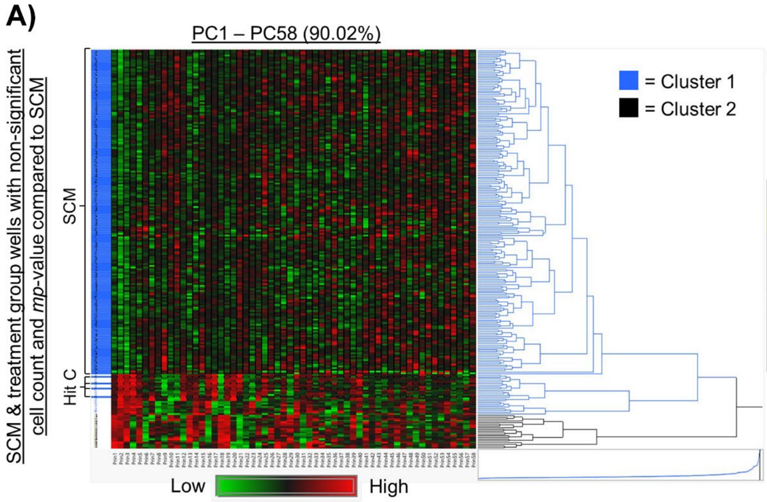


**B)**

| | Cell Count Statistical Comparisons | Morphology Statistical Comparisons |
|---|---|---|
| **Treatment** | **SCM adjusted *p*-value** | **SCM *mp*-value** |
| -TEI-L-A (Hit C) | 1 | 0.2 |
| -TE----- | 1 | 0.171 |
| -T---L-A | 1 | 0.165 |
| -TE----A | 0.725872838 | 0.093 |
| -TE--L-A | 0.689338684 | 0.062 |
| -T-I-LSA | 1 | 0.052 |

**Figure S3: Hierarchical clustering and mp-value testing identify Hit J as CDM formulation most similar to SCM.** A) Hierarchical clustering using 58 principal components that represent 90% of the variance in the data of all SCM control wells and treatment groups with non-significant cell counts and mp-values compared to SCM. B) Table showing ranking by mp-value of treatment groups that cluster with SCM. Letters represent growth factors included in the treatment group formulations.

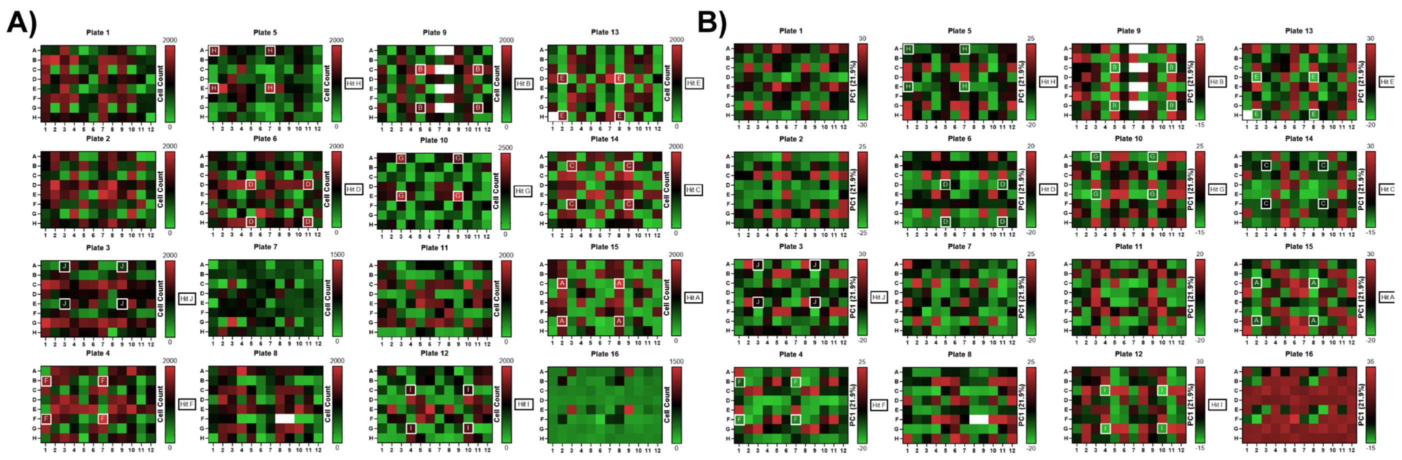

**Figure S4: Visual appraisal of exploratory HTS heatmaps serves as quality assessment for hit identification.** Heatmaps representing A) cell count and B) PC1. Plates were organized into randomized quadrants of unique conditions, where each condition was repeated across four replicate wells per plate.

A)

| Plate 1: AD-MSC RB98 | | |
|---|---|---|
| Treatment | CDBM *mp* -value | SCM *mp* -value |
| CDBM | 1 | 0 |
| SCM | 0 | 0.997 |
| Hit A | 0 | 0 |
| Hit B | 0.001 | 0 |
| Hit C | 0 | 0 |
| Hit D | 0 | 0 |
| Hit E | 0 | 0 |
| Hit F | 0 | 0 |
| Hit G | 0 | 0.001 |
| Hit H | 0 | 0.001 |
| Hit I | 0 | 0 |
| Hit J | 0.002 | 0 |

| Plate 2: BM-MSC RB71 | | |
|---|---|---|
| Treatment | CDBM *mp* -value | SCM *mp* -value |
| CDBM | 1 | 0 |
| SCM | 0 | 0.998 |
| Hit A | 0 | 0 |
| Hit B | 0 | 0.001 |
| Hit C | 0 | 0 |
| Hit D | 0 | 0 |
| Hit E | 0.001 | 0.001 |
| Hit F | 0 | 0 |
| Hit G | 0 | 0 |
| Hit H | 0 | 0 |
| Hit I | 0.001 | 0.001 |
| Hit J | 0 | 0 |

| Plate 3: BM-MSC RB115 | | |
|---|---|---|
| Treatment | CDBM *mp* -value | SCM *mp* -value |
| CDBM | 1 | 0 |
| SCM | 0 | 0.999 |
| Hit A | 0 | 0.001 |
| Hit B | 0.001 | 0 |
| Hit C | 0.001 | 0.001 |
| Hit D | 0 | 0 |
| Hit E | 0 | 0 |
| Hit F | 0 | 0 |
| Hit G | 0 | 0 |
| Hit H | 0.001 | 0.001 |
| Hit I | 0.001 | 0 |
| Hit J | 0 | 0 |

B)

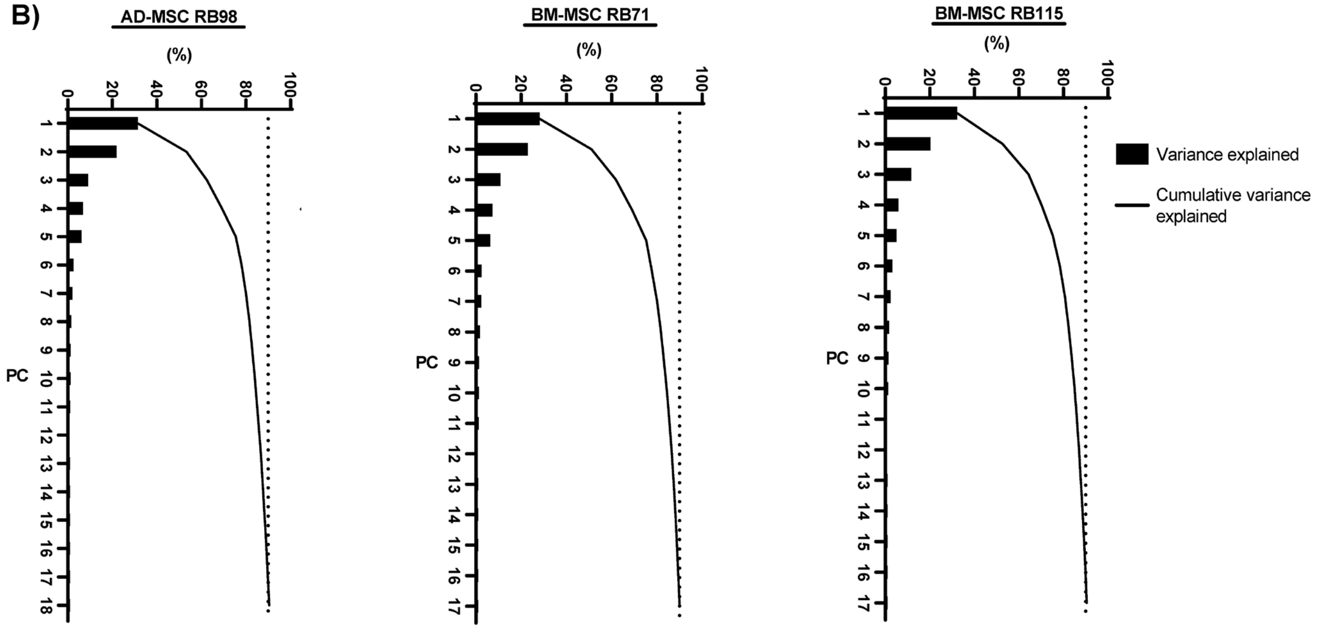


**Figure S5: Validation HTS mp-value testing highlights phenotypic differences between hits and controls using 18 principal components that represent 90% of the variance in the data.** A) mp-values for phenotypic statistical comparisons between each hit and on-plate controls for each donor. B) Validation HTS PCA cumulative variance plot showing number of PC’s that represent 90% of the variance in the data for each donor.

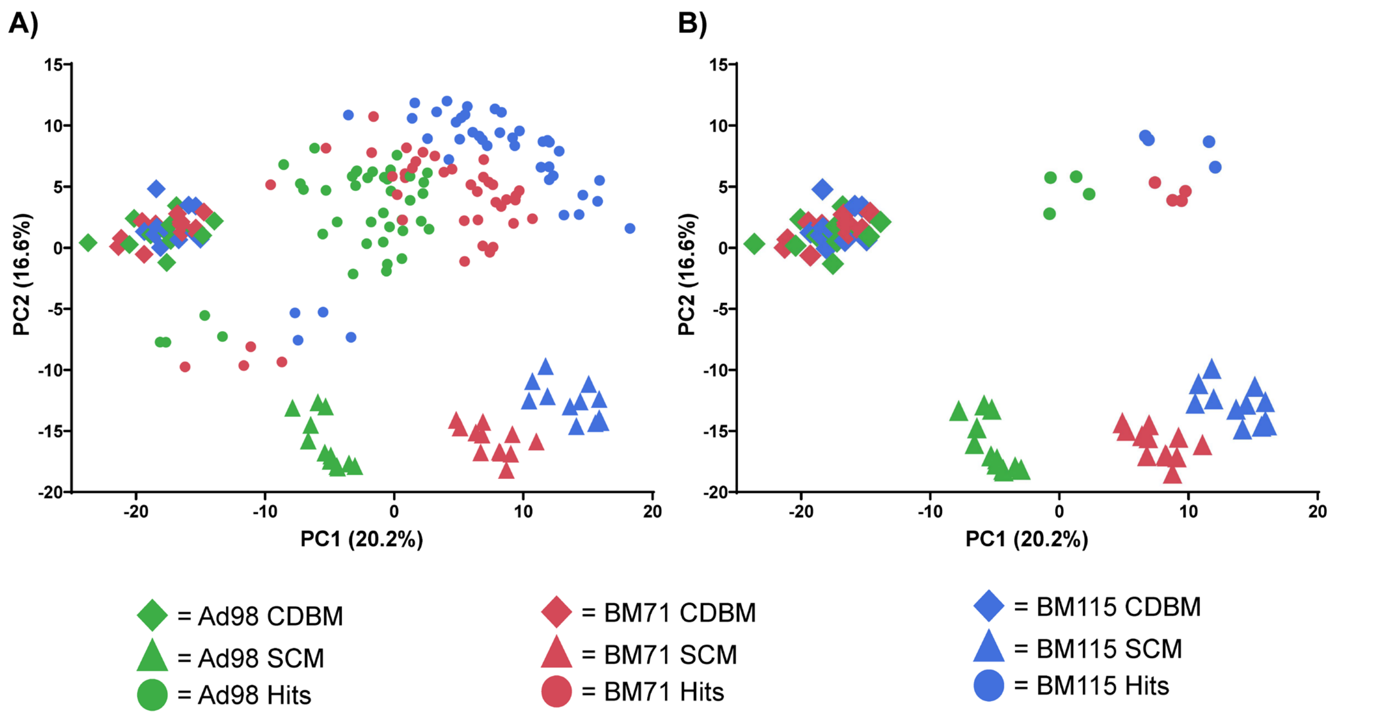


**Figure S6: Heterogeneity of morphological responses to different media types persists across multiple MSC donors.** PCA plots representing morphological differences across all donors cultured in different media types during validation screen: A) all hits, SCM, CDBM. B) Hit G, SCM, CDBM. Morphological data normalized to CDBM.

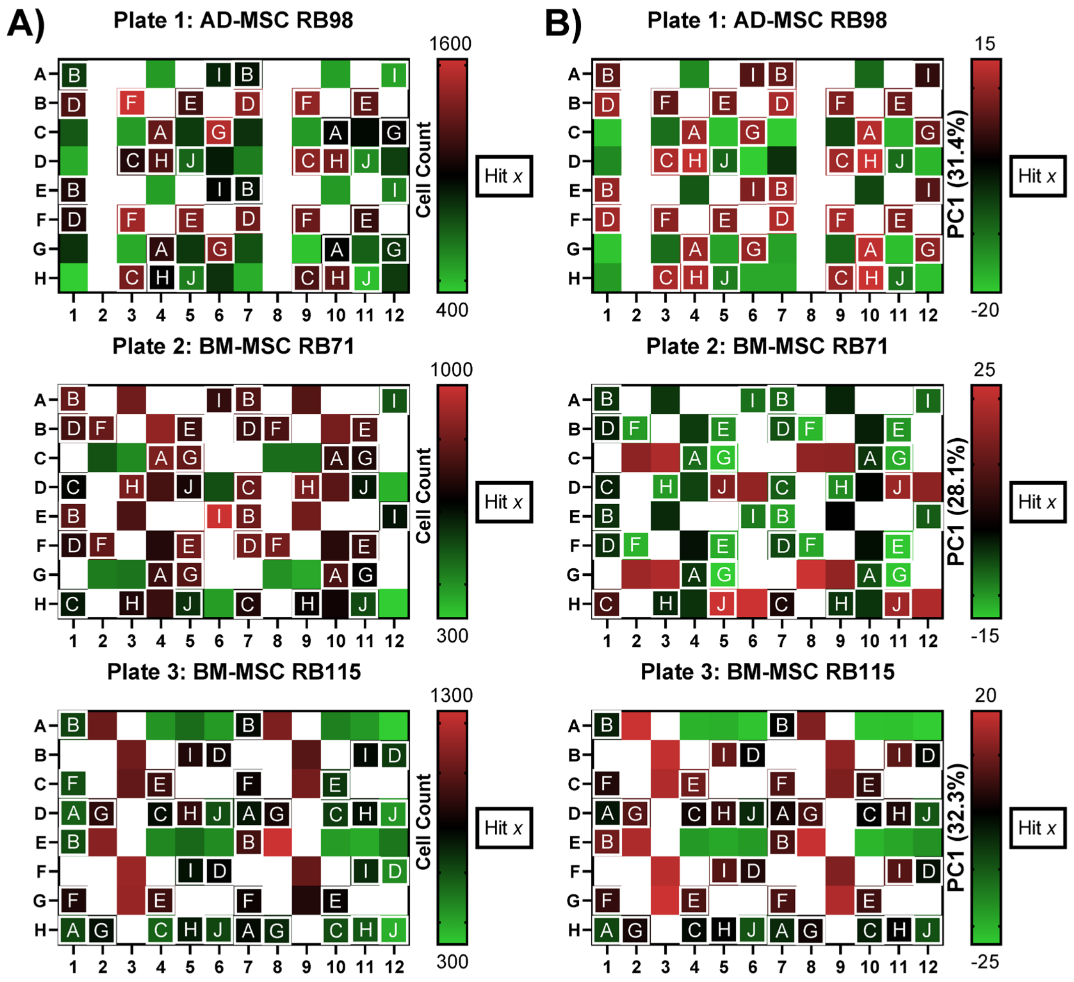


**Figure S7: Visual quality assessment of validation HTS heatmaps.** Heatmaps representing A) cell count and B) PC1. Plates were organized into randomized quadrants of unique conditions, where each condition was repeated across four replicate wells per plate.

A)

| | Morphology Statistical Comparison | | |
|---|---|---|---|
| **Treatment** | **SCM *mp* -value** | **CDBM *mp* -value** | **+CTL *mp* -value** |
| Hit[Low] | 0.03 | 0.003 | 0.854 |
| Hit[Med] | 0.001 | 0 | 0.143 |

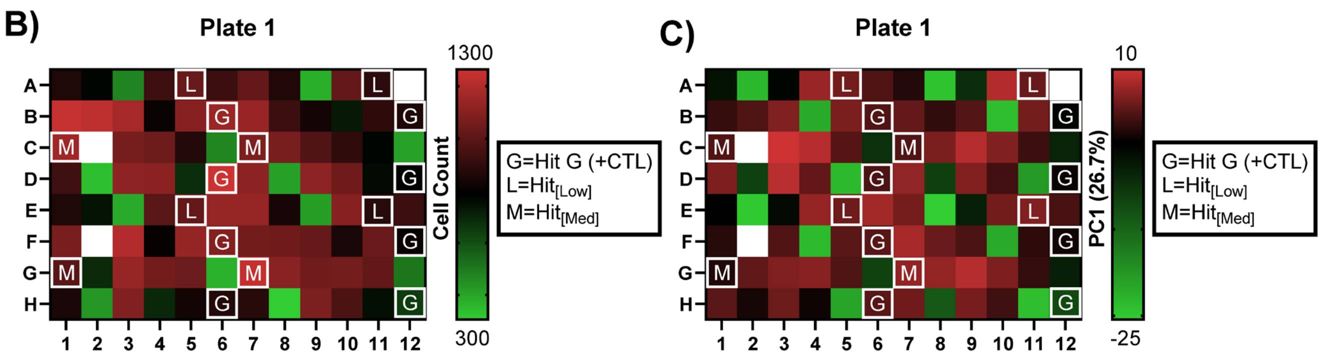


**Figure S8: Refinement HTS mp-value testing highlights phenotypic differences between hits and negative controls but similar morphology between hits and +CTL, while heatmaps support replicable biological phenomena of hits.** A) mp-values for phenotypic statistical comparisons between each hit and on-plate controls for each donor. Visual refinement HTS quality assessment heatmaps for A) cell count and B) PC1. Plates were organized into randomized quadrants of unique conditions, where each condition was repeated across four replicate wells per plate.

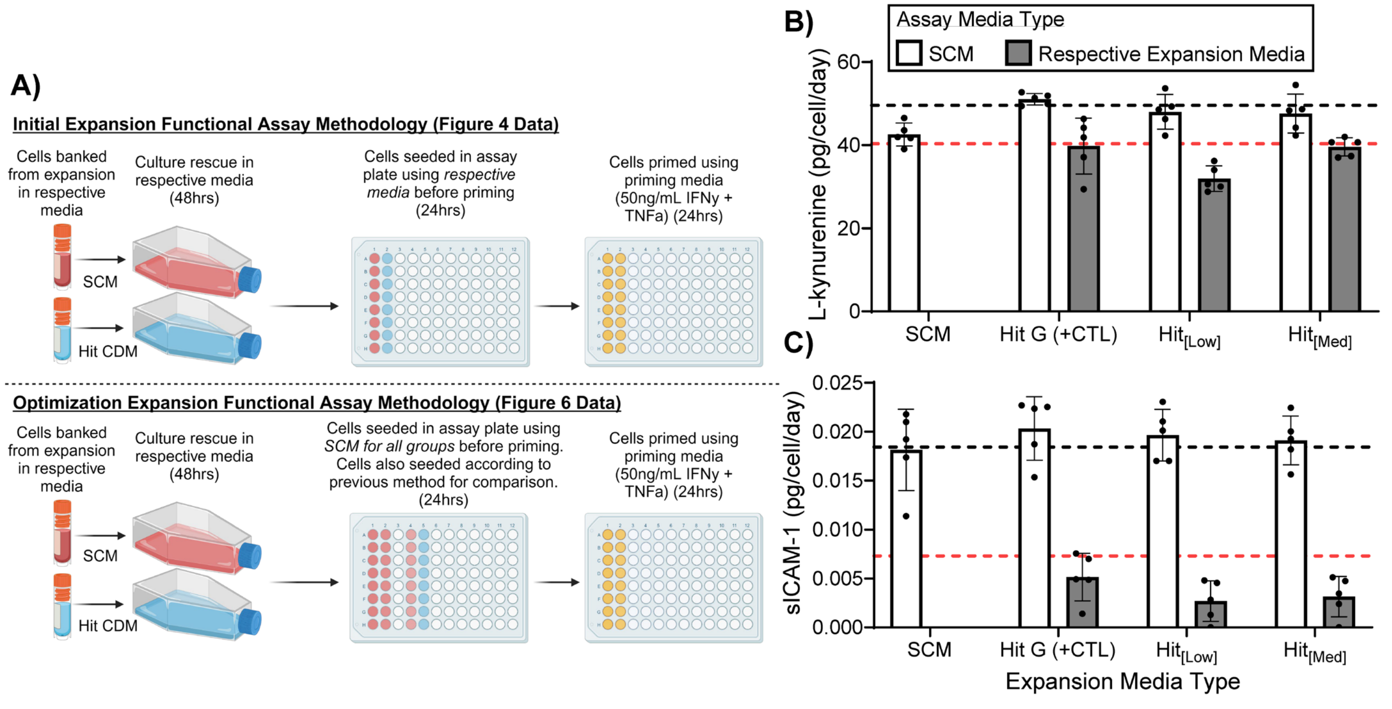


**Figure S9: Functional assay methodology impacts MSC response to priming.** A) Schematic representing differences in functional assay methodology tested after refinement screen expansion. B) IDO activity versus expansion media type and media type used for seeding in assay plate. Black dashed line = L-kynurenine (pg/cell/day) after P1 of initial expansion for MSCs grown in SCM (Fig. 4E). Red dashed line = L-kynurenine (pg/cell/day) after P1 of initial expansion for MSCs grown in Hit G (+CTL) (Fig. 4E). C) sICAM-1 secretion versus expansion media type and media type used for seeding in assay plate. Black dashed line = sICAM-1 (pg/cell/day) after P1 of initial expansion for MSCs grown in SCM (Fig. 4E). Red dashed line = sICAM-1 (pg/cell/day) after P1 of initial expansion for MSCs grown in Hit G (+CTL) (Fig. 4E).
