## Supplementary material for "High Throughput Morphological Screening Identifies Chemically Defined Media for Mesenchymal Stromal Cells that Enhances Proliferation and Supports Maintenance of Immunomodulatory Function": S3_1.profile-QCFull_v1.1.py

```
#!/usr/bin/env python
# coding: utf-8

# # Apply an image-based profiling pipeline using pycytominer
# 
# As described fully in [Caicedo et al. 2017](https://doi.org/10.1038/nmeth.4397), an image-based profiling pipeline consists of three core steps:
# 
# 1. Aggregation
# 2. Normalization
# 3. Feature selection
# 
# [Pycytominer](https://github.com/cytomining/pycytominer) is a python package, built on top of pandas, that facilitates all of these steps and more.
# 
# ### Data levels
# 
# The concept of "data levels" is important to understand when implementing an image-based profiling pipeline.
# 
# | Data | Level |
# | :---- | :---- |
# | Images | Level 1 |
# | Single cell profiles (SQLite) | Level 2 |
# | Aggregated profiles with metadata information | Level 3 |
# | Normalized aggregated profiles | Level 4a |
# | Normalized and feature selected profiles | Level 4b |
# | Consensus profiles | Level 5 |
# 
# 
# ### Script Workflow & Checks:
# 1. Initialize data:
# 
#     A. Check for NaN values in platemap_df
#     
#     B. Check that shape of cells, nucleus, and cytoplasm is equal for each plate
#     
#     C. Merge cells, nucleus, and cytoplasm.csvs together instead of concatenate
#     
#     D. Feature Engineering
#     
#     E. Check for NaN values in 'Metadata_Plate' & 'Metadata_Well' in full_data once merged together
#     
#     
# 2. Single Cell Aggregation
# 
#     A. Calculate well medians
#     
#     B. Merge cell count data (N dataframe) to medians dataframe, output .csv
#     
#     C. Check datatype of 'Metadata_Plate' and 'Metadata_Well' columns
#     
#     D. Annotate medians dataframe with platemap data
#     
# 
# 3. Normalize using RobustMAD Method
# 
#     A. Normalization done on a per-plate basis, output .csv
#     
#     
# 4. Feature Selection
# 
#     A. Perform correlation_threshold, with threshold value = 0.9, output .csv

# In[ ]:


import pathlib
import pandas as pd
import numpy as np

from pycytominer import aggregate, annotate, normalize, feature_select, consensus
pd.options.mode.chained_assignment = None


# ### Step 0 - Initialize data

# In[ ]:


# Load all .csv files
platemap = r"Conditions.csv"
cells_csv_path = r"=Cells.csv"
nuclei_csv_path = r"FilteredNuclei.csv"
cytoplasm_csv_path = r"Cytoplasm.csv"

platemap_df = pd.read_csv(platemap)
cells_ori = pd.read_csv(cells_csv_path)
nuclei_ori = pd.read_csv(nuclei_csv_path)
cytoplasm_ori = pd.read_csv(cytoplasm_csv_path)

print("Original Cells.csv size (row,col): ", cells_ori.shape)
print("Original Nuclei.csv size (row,col): ", nuclei_ori.shape)
print("Original Cytoplasm.csv size (row,col): ", cytoplasm_ori.shape)
print()


# In[ ]:


# Delete duplicate columns (by column name)
cells = cells_ori.loc[:,cells_ori.columns.str.find('.') < 0]
nuclei = nuclei_ori.loc[:,nuclei_ori.columns.str.find('.') < 0]
cytoplasm = cytoplasm_ori.loc[:,cytoplasm_ori.columns.str.find('.') < 0]

print("Cells.csv size after removing duplicate cols (row,col): ", cells.shape)
print("Nuclei.csv size after removing duplicate cols (row,col): ", nuclei.shape)
print("Cytoplasm.csv size after removing duplicate cols (row,col): ", cytoplasm.shape)
print()


# In[ ]:


# Check to see how Cell data looks
print("Cell data (rows, columns):", cells.shape)
print()
cells.head(5)


# In[ ]:


# Check to see how platemap/conditions file looks
print("Conditions data (rows, columns):", platemap_df.shape)
print("Sum of NaN values in conditions data: ", platemap_df.isnull().sum().sum())
print()
platemap_df.head(6)


# In[ ]:


# Scan through column headers in .csvs until you reach "AreaShape_Area", then append prefix to every column thereafter
x = cells.columns.get_loc("AreaShape_Area")
new_cell_names = [(i,'Cells_'+i) for i in cells.iloc[:, x:].columns.values]
cells.rename(columns = dict(new_cell_names), inplace=True)

x = nuclei.columns.get_loc("AreaShape_Area")
new_nuclei_names = [(i,'Nuclei_'+i) for i in nuclei.iloc[:, x:].columns.values]
nuclei.rename(columns = dict(new_nuclei_names), inplace=True)

x = cytoplasm.columns.get_loc("AreaShape_Area")
new_cytoplasm_names = [(i,'Cytoplasm_'+i) for i in cytoplasm.iloc[:, x:].columns.values]
cytoplasm.rename(columns = dict(new_cytoplasm_names), inplace=True)

cytoplasm.head(5)


# In[ ]:


cells_plates = {}
# Iterate over each unique plate in the csv
print("Plates Identified in Cells.csv: ")
for plate_num in cells['Metadata_Plate'].unique():
    cells_plates[plate_num] = cells[cells['Metadata_Plate'] == plate_num]
    print(f'Identified plate {plate_num} with shape {cells_plates[plate_num].shape}')
print()

print("Plates Identified in Nuclei.csv: ")
nuclei_plates = {}
# Iterate over each unique plate in the csv
for plate_num in nuclei['Metadata_Plate'].unique():
    nuclei_plates[plate_num] = nuclei[nuclei['Metadata_Plate'] == plate_num]
    print(f'Identified plate {plate_num} with shape {nuclei_plates[plate_num].shape}')
print()

print("Plates Identified in Cytoplasm.csv: ")
cyto_plates = {}
# Iterate over each unique plate in the csv
for plate_num in cytoplasm['Metadata_Plate'].unique():
    cyto_plates[plate_num] = cytoplasm[cytoplasm['Metadata_Plate'] == plate_num]
    print(f'Identified plate {plate_num} with shape {cyto_plates[plate_num].shape}')
print()


# In[ ]:


# Get cell number from cells.csv with 'Metadata_Plate' & 'Metadata_Well' information conserved
N = cells.groupby(['Metadata_Plate','Metadata_Well'],as_index=False).size()
N['Plate_Well'] = N['Metadata_Plate'].astype(str) + N['Metadata_Well']
print("Shape of cell count dataframe: ", N.shape)


# In[ ]:


# Merge Cells, Nuclei, Cytoplasm data together, delete duplicated 'Metadata' columns

# Make 'Metadata_Signature' for each .csv from ImageNumber, Site, ObjectNumber, Plate, and Well:
cells['Signature'] = cells['ImageNumber'].astype(str) + cells['Metadata_Site'].astype(str) + cells['ObjectNumber'].astype(str) + cells['Metadata_Plate'].astype(str) + cells['Metadata_Well']
nuclei['Signature'] = nuclei['ImageNumber'].astype(str) + nuclei['Metadata_Site'].astype(str) + nuclei['ObjectNumber'].astype(str) + nuclei['Metadata_Plate'].astype(str) + nuclei['Metadata_Well']
cytoplasm['Signature'] = cytoplasm['ImageNumber'].astype(str) + cytoplasm['Metadata_Site'].astype(str) + cytoplasm['ObjectNumber'].astype(str) + cytoplasm['Metadata_Plate'].astype(str) + cytoplasm['Metadata_Well']

# First merge cells and nuclei:
half_data = pd.merge(cells,nuclei,on='Signature',suffixes=('','_remove'))
print("Shape of cells+nucleus dataframe (including duplicate cols): ", half_data.shape)

# Remove duplicate columns:
half_data.drop([i for i in half_data.columns if 'remove' in i], axis=1, inplace=True)
print("Shape of cells+nucleus dataframe after removing duplicate cols: ", half_data.shape)

# Merge half_data and cytoplasm to make 'full_data':
full_data = pd.merge(half_data,cytoplasm,on='Signature',suffixes=('','_remove'))
print("Shape of cells+nucleus+cyto dataframe (including duplicate cols): ", full_data.shape)
print()

# Remove duplicate columns:
full_data.drop([i for i in full_data.columns if 'remove' in i], axis=1, inplace=True)
print("Shape of full dataframe after removing duplicate cols: ", full_data.shape)
print()


# In[ ]:


# Feature Engineering - Addition of new data columns based on calculation from existing data
full_data['Cells_NC_Ratio'] = full_data['Nuclei_AreaShape_Area']/full_data['Cells_AreaShape_Area']
full_data.info()
print("Full data (rows, cols) after adding NC Ratio: ", full_data.shape)


# In[ ]:


full_data['Cells_Aspect_Ratio_Cell'] = full_data['Cells_AreaShape_MajorAxisLength']/full_data['Cells_AreaShape_MinorAxisLength']


# In[ ]:


full_data['Cells_Aspect_Ratio_Nucleus'] = full_data['Nuclei_AreaShape_MajorAxisLength']/full_data['Nuclei_AreaShape_MinorAxisLength']


# In[ ]:


full_data['Cells_PerimeterArea_Ratio_Cell'] = full_data['Cells_AreaShape_Perimeter']/full_data['Cells_AreaShape_Area']


# In[ ]:


full_data['Cells_PerimeterArea_Ratio_Nucleus'] = full_data['Nuclei_AreaShape_Perimeter']/full_data['Nuclei_AreaShape_Area']
full_data.info()
full_data.head(10)
print("Full data (rows,cols) after adding all additional features: ", full_data.shape)


# In[ ]:


# Filter data based on NC Ratio, Nucleus Solidity, & Cell Solidity

# Find outliers based on NC Ratio >= 0.75
full_data.groupby(pd.cut(full_data['Cells_NC_Ratio'], np.arange(0, 1.1, 0.1)))['Metadata_Well'].count()
outlier_objects = full_data[full_data['Cells_NC_Ratio'] >= 0.75][['Metadata_Well', 'ObjectNumber', 'Cells_NC_Ratio']]
outlier_objects.to_csv('outlier_images.csv',index=False)


# In[ ]:


# Drop NC Ratio outliers
full_data.drop(full_data[full_data.Cells_NC_Ratio >= 0.75].index, inplace=True)
print("Full data (row,col) after filtering based on NC Ratio: ", full_data.shape)

# Drop Nucleus Solidity outliers
full_data.drop(full_data[full_data.Nuclei_AreaShape_Solidity > 1].index, inplace=True)
print("Full data (row,col) after filtering based on Nucleus Solidity: ", full_data.shape)

# Drop Cell Solidity outliers
full_data.drop(full_data[full_data.Cells_AreaShape_Solidity > 1].index, inplace=True)
print("Full data (row,col) after filtering based on Cell Solidity: ", full_data.shape)
print()


# In[ ]:


# Check for NaN values in full_data for 'Metadata_Plate' & 'Metadata_Well'
x = full_data['Metadata_Plate'].isnull().sum()
print("sum of NaN values in Metadata_Plate: ", x)

x = full_data['Metadata_Well'].isnull().sum()
print("sum of NaN values in Metadata_Well: ", x)
print()

full_data = full_data.rename(columns={'ImageNumber': 'Metadata_ImageNumber', 'ObjectNumber': 'Metadata_ObjectNumber'})


# ### Step 1 - Single cell aggregation
# 
# In this step, **level 2 profiles** (single cells) are processed to **level 3 profiles** (well-level profiles).

# In[ ]:


# Perform the aggregation - output well level profiles
medians = aggregate(
    full_data,
    strata=["Metadata_Plate", "Metadata_Well"],
    features="infer",
    operation="median",
    subset_data_df="none",
)
print("Median profile size (row,col):", medians.shape)

# Merge medians and cell count data: 
medians['Plate_Well'] = medians['Metadata_Plate'].astype(str) + medians['Metadata_Well']
medians = pd.merge(medians,N,on='Plate_Well',suffixes=('','_remove'))

# Remove duplicate columns: 
medians.drop([i for i in medians.columns if 'remove' in i], axis=1, inplace=True)
print("Shape of medians and cell count after removing duplicate cols: ", medians.shape)
cols = ['size','Plate_Well','Metadata_Plate','Metadata_Well']
medians = medians[cols + [c for c in medians.columns if c not in cols]]
medians.head(12)


# ## Step 2 - Annotate wells using the platemap file
# 
# In this step, **level 3 profiles** (well-level profiles) are annotated with platemap metadata.

# In[ ]:


# Annotate medians dataframe with platemap/conditions file

# Check data type of 'Metadata_Plate' & 'Metadata_Well':
print("Data type of Metadata_Plate: ", medians['Metadata_Plate'].dtypes)
print("Data type of Metadata_Well: ", medians['Metadata_Well'].dtypes)

# Merge platemap and medians dataframes: 
medians_annotated = pd.merge(medians,platemap_df,left_on='Plate_Well',right_on='Metadata_Plate_Well',suffixes=('','_remove'))
print("medians_annotated size after merge: ", medians_annotated.shape)

# Drop duplicate columns that are made when appending platemap/conditions dataframe
medians_annotated.drop([i for i in medians_annotated.columns if 'remove' in i], axis=1, inplace=True)
medians_annotated.rename(columns = {"size": "Metadata_size"}, inplace = True)
print("medians_annotated size after merge and removing duplicate cols: ", medians_annotated.shape)

print("Writing medians.csv: ")
medians_annotated.to_csv("medians.csv", index=False)
medians_annotated.head(5)


# ## Step 3 - Normalize well-level profiles
# 
# In this step, **level 3 profiles** (well-level profiles) are normalized to form **level 4a profiles**.

# In[ ]:


# Create a dictionary (plates) which looks like this : {'1': plate1, '2': plate2, etc.}
# plate1 and plate2 contain rows corresponding to plates 1 and 2 only respectively
plates = {}

# Iterate over each unique plate in the csv
for plate_num in medians_annotated['Metadata_Plate'].unique():
    plates[plate_num] = medians_annotated[medians_annotated['Metadata_Plate'] == plate_num]
    print(f'Identified plate {plate_num} with shape {plates[plate_num].shape}')


# In[ ]:


# Normalize profiles by plate
medians_normalized = []
for plate_num, plate in plates.items():
    normalized_plate = normalize(
        plates[plate_num],
        features="infer",
        meta_features="infer",
        samples="Metadata_Treatment == '-CTL'",
        method="mad_robustize",
    )
    medians_normalized.append(normalized_plate)
    
medians_normalized = pd.concat(medians_normalized)
medians_normalized.head(12)
print("Writing medians_normalized.csv: ")
medians_normalized.to_csv("medians_normalized.csv", index=False)


# ## Step 4 - Apply feature selection
# 
# In this step, we apply a series of feature selection steps to **level 4a profiles** (normalized well-level profiles) to form **level 4b profiles**.

# In[ ]:


# Apply feature selection to all data
feature_select_opts = [
    "variance_threshold",
    "drop_na_columns",
    "correlation_threshold",
    "blocklist",
    "drop_outliers"
]

normalized_features = feature_select(
    profiles=medians_normalized,
    features="infer",
    samples="all",
    operation=["correlation_threshold"],
    corr_threshold=0.9,
    freq_cut=0.05,
    unique_cut=0.1,
    noise_removal_perturb_groups= "Metadata_outliers",
    noise_removal_stdev_cutoff= 3,
)
normalized_features.head(5)
print("Writing final_data_for_jmp.csv: ")
normalized_features.to_csv("final_data_for_jmp.csv", index=False)


# ## Step 5 - Form consensus signatures
# 
# In this step, we collapse replicates (**level 4 profiles**) into a single profile.
# This forms **level 5 profiles**.

# In[ ]:


# Generate consensus profiles
#consensus_file = pathlib.Path(f"{output_dir}/{plate_id}_consensus.csv.gz")

#consensus(
    #profiles=feature_select_df,
    #replicate_columns=["Metadata_clone_number", "Metadata_treatment"],
    #operation="modz",
    #features="infer",
    #output_file=consensus_file,
    #modz_method="spearman",
    #modz_min_weight=0.01,
    #modz_precision=4,
    #compression="gzip",
#)


# In[ ]:


# Read in and preview what was output in the previous step 
#consensus_df = pd.read_csv(consensus_file)

#print(consensus_df.shape)
#consensus_df.head(3)
```
