## Supplementary material for "High Throughput Morphological Screening Identifies Chemically Defined Media for Mesenchymal Stromal Cells that Enhances Proliferation and Supports Maintenance of Immunomodulatory Function": S4_cytominer-eval_mpValue.py

```
#!/usr/bin/env python
# coding: utf-8

# In[1]:


import numpy as np
import pandas as pd
from sklearn import preprocessing

from cytominer_eval import evaluate
from pycytominer.cyto_utils import infer_cp_features


# In[2]:


#Load Data
df = pd.read_csv("final_data_for_jmp.csv")
print(df.shape)
df = df.dropna(axis=0)
df.head(10)


# In[3]:


#Set Feature Information
features = infer_cp_features(df)
meta_features = infer_cp_features(df,metadata=True)

meta_features


# In[4]:


#define control perturbation group
control_perts = df.query("Metadata_Treatment == 'GM-MSC'").Metadata_Treatment.unique().tolist()
control_perts
print(control_perts)


# In[5]:


mp_values = []
    
for Plate in df.Metadata_Plate.unique():
    Plate_df = df.query("Metadata_Plate == @Plate")
    print(Plate_df.head())
    
    for num_perm in [1000]:
    
            mp_results = evaluate(
                profiles=Plate_df,
                features=features,
                meta_features=meta_features,
                replicate_groups="Metadata_Treatment",
                operation="mp_value",
                similarity_metric="pearson",
                mp_value_params={"rescale_pca": True, "nb_permutations": num_perm},
                grit_control_perts=control_perts
                ).assign(Plate=Plate, permutations=num_perm)
        
            mp_values.append(mp_results)


# In[6]:


mp_values = pd.concat(mp_values).reset_index(drop=True)

sig_05_mp_values = mp_values.loc[mp_values["mp_value"] <= 0.05]
sig_005_mp_values = mp_values.loc[mp_values["mp_value"] <= 0.005]
sig_001_mp_values = mp_values.loc[mp_values["mp_value"] <= 0.001]
sig_0001_mp_values = mp_values.loc[mp_values["mp_value"] <= 0.0001]

mp_values.to_csv("mp_values.csv", index=False)
sig_05_mp_values.to_csv("sig_05_mp_values.csv", index=False)
sig_005_mp_values.to_csv("sig_005_mp_values.csv", index=False)
sig_001_mp_values.to_csv("sig_001_mp_values.csv", index=False)
sig_0001_mp_values.to_csv("sig_0001_mp_values.csv", index=False)

print(mp_values.shape)
print(sig_05_mp_values.shape)
print(sig_005_mp_values.shape)
print(sig_001_mp_values.shape)
print(sig_0001_mp_values.shape)
sig_05_mp_values.head()


# In[7]:


#Collect non-significant groups
nonsig_mp_values = mp_values.loc[mp_values["mp_value"] > 0.05]

nonsig_mp_values.to_csv("nonsig_mp_values.csv", index=False)

print(nonsig_mp_values.shape)
```
