## Supplementary material for "High Throughput Morphological Screening Identifies Chemically Defined Media for Mesenchymal Stromal Cells that Enhances Proliferation and Supports Maintenance of Immunomodulatory Function": S5_representative_Cell_images.py

```
#!/usr/bin/env python
# coding: utf-8

# # Apply an image-based profiling pipeline using pycytominer
# 
# As described fully in [Caicedo et al. 2017](https://doi.org/10.1038/nmeth.4397), an image-based profiling pipeline consists of three core steps:
# 
# 1. Aggregation
# 2. Normalization
# 3. Feature selection
# 
# [Pycytominer](https://github.com/cytomining/pycytominer) is a python package, built on top of pandas, that facilitates all of these steps and more.
# 
# ### Data levels
# 
# The concept of "data levels" is important to understand when implementing an image-based profiling pipeline.
# 
# | Data | Level |
# | :---- | :---- |
# | Images | Level 1 |
# | Single cell profiles (SQLite) | Level 2 |
# | Aggregated profiles with metadata information | Level 3 |
# | Normalized aggregated profiles | Level 4a |
# | Normalized and feature selected profiles | Level 4b |
# | Consensus profiles | Level 5 |
# 
# 
# ### Script Workflow & Checks:
# 1. Initialize data:
# 
#     A. Check for NaN values in platemap_df
#     
#     B. Check that shape of cells, nucleus, and cytoplasm is equal for each plate
#     
#     C. Merge cells, nucleus, and cytoplasm.csvs together instead of concatenate
#     
#     D. Feature Engineering
#     
#     E. Check for NaN values in 'Metadata_Plate' & 'Metadata_Well' in full_data once merged together
#     
#     
# 2. Single Cell Aggregation
# 
#     A. Calculate well medians
#     
#     B. Merge cell count data (N dataframe) to medians dataframe, output .csv
#     
#     C. Check datatype of 'Metadata_Plate' and 'Metadata_Well' columns
#     
#     D. Annotate medians dataframe with platemap data
#     
# 
# 3. Normalize using RobustMAD Method
# 
#     A. Normalization done on a per-plate basis, output .csv
#     
#     
# 4. Feature Selection
# 
#     A. Perform correlation_threshold, with threshold value = 0.9, output .csv

# In[92]:


import pathlib
import pandas as pd
import numpy as np

from pycytominer import aggregate, annotate, normalize, feature_select, consensus
pd.options.mode.chained_assignment = None


# ### Step 0 - Initialize single-cell data

# In[93]:


# Load all .csv files
cells_csv_path = r"Cells.csv"
nuclei_csv_path = r"FilteredNuclei.csv"
medians_csv = r"medians.csv"

cells_ori = pd.read_csv(cells_csv_path)
nuclei_ori = pd.read_csv(nuclei_csv_path)
medians = pd.read_csv(medians_csv)

print("Original Cells.csv size (row,col): ", cells_ori.shape)
print("Original Nuclei.csv size (row,col): ", nuclei_ori.shape)
print("Original medians.csv size (row,col): ", medians.shape)
print()


# In[94]:


# Delete duplicate columns (by column name)
cells = cells_ori.loc[:,cells_ori.columns.str.find('.') < 0]
nuclei = nuclei_ori.loc[:,nuclei_ori.columns.str.find('.') < 0]

print("Cells.csv size after removing duplicate cols (row,col): ", cells.shape)
print("Nuclei.csv size after removing duplicate cols (row,col): ", nuclei.shape)
print()


# In[95]:


# Check to see how Cell data looks
print("Cell data (rows, columns):", cells.shape)
print()
cells.head(5)


# In[97]:


# Scan through column headers in .csvs until you reach "AreaShape_Area", then append prefix to every column thereafter
x = cells.columns.get_loc("AreaShape_Area")
new_cell_names = [(i,'Cells_'+i) for i in cells.iloc[:, x:].columns.values]
cells.rename(columns = dict(new_cell_names), inplace=True)

x = nuclei.columns.get_loc("AreaShape_Area")
new_nuclei_names = [(i,'Nuclei_'+i) for i in nuclei.iloc[:, x:].columns.values]
nuclei.rename(columns = dict(new_nuclei_names), inplace=True)

cells.head(5)


# In[98]:


cells_plates = {}
# Iterate over each unique plate in the csv
print("Plates Identified in Cells.csv: ")
for plate_num in cells['Metadata_Plate'].unique():
    cells_plates[plate_num] = cells[cells['Metadata_Plate'] == plate_num]
    print(f'Identified plate {plate_num} with shape {cells_plates[plate_num].shape}')
print()

print("Plates Identified in Nuclei.csv: ")
nuclei_plates = {}
# Iterate over each unique plate in the csv
for plate_num in nuclei['Metadata_Plate'].unique():
    nuclei_plates[plate_num] = nuclei[nuclei['Metadata_Plate'] == plate_num]
    print(f'Identified plate {plate_num} with shape {nuclei_plates[plate_num].shape}')
print()


# In[100]:


# Merge Cells and Nuclei data together, delete duplicated 'Metadata' columns

# Make 'Metadata_Signature' for each .csv from ImageNumber, Site, ObjectNumber, Plate, and Well:
cells['Signature'] = cells['ImageNumber'].astype(str) + cells['Metadata_Site'].astype(str) + cells['ObjectNumber'].astype(str) + cells['Metadata_Plate'].astype(str) + cells['Metadata_Well']
nuclei['Signature'] = nuclei['ImageNumber'].astype(str) + nuclei['Metadata_Site'].astype(str) + nuclei['ObjectNumber'].astype(str) + nuclei['Metadata_Plate'].astype(str) + nuclei['Metadata_Well']

# Merge cells and nuclei:
full_data = pd.merge(cells,nuclei,on='Signature',suffixes=('','_remove'))
print("Shape of cells+nucleus dataframe (including duplicate cols): ", full_data.shape)

# Remove duplicate columns:
full_data.drop([i for i in full_data.columns if 'remove' in i], axis=1, inplace=True)
print("Shape of full dataframe after removing duplicate cols: ", full_data.shape)
print()


# In[101]:


# Feature Engineering - Addition of new data columns based on calculation from existing data
full_data['Cells_NC_Ratio'] = full_data['Nuclei_AreaShape_Area']/full_data['Cells_AreaShape_Area']
full_data.info()
print("Full data (rows, cols) after adding NC Ratio: ", full_data.shape)


# In[102]:


full_data['Cells_Aspect_Ratio_Cell'] = full_data['Cells_AreaShape_MajorAxisLength']/full_data['Cells_AreaShape_MinorAxisLength']


# In[103]:


full_data['Cells_Aspect_Ratio_Nucleus'] = full_data['Nuclei_AreaShape_MajorAxisLength']/full_data['Nuclei_AreaShape_MinorAxisLength']


# In[104]:


full_data['Cells_PerimeterArea_Ratio_Cell'] = full_data['Cells_AreaShape_Perimeter']/full_data['Cells_AreaShape_Area']


# In[105]:


full_data['Cells_PerimeterArea_Ratio_Nucleus'] = full_data['Nuclei_AreaShape_Perimeter']/full_data['Nuclei_AreaShape_Area']
full_data.info()
full_data.head(10)
print("Full data (rows,cols) after adding all additional features: ", full_data.shape)


# In[106]:


# Filter data based on NC Ratio, Nucleus Solidity, & Cell Solidity

# Find outliers based on NC Ratio >= 0.75
full_data.groupby(pd.cut(full_data['Cells_NC_Ratio'], np.arange(0, 1.1, 0.1)))['Metadata_Well'].count()
outlier_objects = full_data[full_data['Cells_NC_Ratio'] >= 0.75][['Metadata_Well', 'ObjectNumber', 'Cells_NC_Ratio']]
outlier_objects.to_csv('outlier_images.csv',index=False)


# In[107]:


# Drop NC Ratio outliers
full_data.drop(full_data[full_data.Cells_NC_Ratio >= 0.75].index, inplace=True)
print("Full data (row,col) after filtering based on NC Ratio: ", full_data.shape)

# Drop Nucleus Solidity outliers
full_data.drop(full_data[full_data.Nuclei_AreaShape_Solidity > 1].index, inplace=True)
print("Full data (row,col) after filtering based on Nucleus Solidity: ", full_data.shape)

# Drop Cell Solidity outliers
full_data.drop(full_data[full_data.Cells_AreaShape_Solidity > 1].index, inplace=True)
print("Full data (row,col) after filtering based on Cell Solidity: ", full_data.shape)
print()


# In[108]:


# Check for NaN values in full_data for 'Metadata_Plate' & 'Metadata_Well'
x = full_data['Metadata_Plate'].isnull().sum()
print("sum of NaN values in Metadata_Plate: ", x)

x = full_data['Metadata_Well'].isnull().sum()
print("sum of NaN values in Metadata_Well: ", x)
print()

full_data = full_data.rename(columns={'ImageNumber': 'Metadata_ImageNumber', 'ObjectNumber': 'Metadata_ObjectNumber'})


# In[109]:


#Define features of interest & columns to keep:
measurement_columns = ['Cells_AreaShape_Area', 'Cells_AreaShape_Compactness', 'Cells_AreaShape_FormFactor', 
                       'Cells_AreaShape_MajorAxisLength','Cells_AreaShape_MeanRadius', 'Cells_AreaShape_MinorAxisLength',
                      'Cells_AreaShape_Perimeter','Cells_AreaShape_Solidity',
                      'Cells_NC_Ratio','Cells_Aspect_Ratio_Cell','Cells_Aspect_Ratio_Nucleus','Cells_PerimeterArea_Ratio_Cell',
                      'Cells_PerimeterArea_Ratio_Nucleus']

prefix_to_keep = 'Metadata_'

columns_to_keep = measurement_columns + [col for col in full_data.columns if col.startswith(prefix_to_keep)]

full_data = full_data[columns_to_keep]

#Add Plate_Well column
full_data['Metadata_Plate_Well'] = full_data['Metadata_Plate'].astype(str) + full_data['Metadata_Well']

print("Filtered full_data size (row,col): ", full_data.shape)
full_data.head(9)


# ### Step 1 - Isolate representative wells using well-median data
# 
# In this step, **level 3 profiles** (well-level profiles) are processed to find which lies closest to the mean-of-medians value for each feature of interest

# In[110]:


#Define features of interest & columns to keep for medians.csv:
columns_to_keep = measurement_columns + [col for col in medians.columns if col.startswith(prefix_to_keep)]

medians = medians[columns_to_keep]
print("Filtered medians.csv size (row,col): ", medians.shape)
medians.head(9)


# In[111]:


# Initialize an empty dictionary to store the results
isolated_wells = {}

for column in measurement_columns:
    # Calculate the mean for each unique 'Treatment' group
    mean_values = medians.groupby('Metadata_Treatment')[column].mean().reset_index()
    
    # Merge the mean values back into the original DataFrame
    medians = pd.merge(medians, mean_values, on='Metadata_Treatment', suffixes=('', '_mean'))
    
    # For each row, Calculate the absolute difference between the measurement and the mean of each treatment group
    medians[column + '_diff'] = abs(medians[column] - medians[column + '_mean'])
    
    # Find the 'Plate_Well' with the minimum absolute difference for each 'Metadata_Treatment' group
    isolated_wells[column] = medians.loc[medians.groupby('Metadata_Treatment')[column + '_diff'].idxmin()]

print("Calculations added to medians.csv size (row,col): ", medians.shape)
medians.head(16)


# In[112]:


# Create a DataFrame that combines all closest 'Plate_Well' results
isolatedWells_df = pd.concat(isolated_wells, axis=0, ignore_index=True)

print("Shape of dataframe with only minimum absolute difference well remaining (row,col): ", isolatedWells_df.shape)
isolatedWells_df.head(300)


# In[113]:


# Count the occurrences of each 'Plate_Well' for each 'Treatment'
count = isolatedWells_df['Metadata_Plate_Well'].value_counts().reset_index()
count.columns = ['Metadata_Plate_Well', 'Count']

# Merge the "count" DataFrame with the "isolatedWells_df" DataFrame
isolatedWells_df = isolatedWells_df.merge(count, on='Metadata_Plate_Well')

# Group the "isolatedWells_df" DataFrame by 'Metadata_Treatment' and find the row with the highest "Count" for each group
most_common_rows = isolatedWells_df.loc[isolatedWells_df.groupby('Metadata_Treatment')['Count'].idxmax()]

# List of column names you want to keep in "isolatedWells_df"
selected_columns = ['Metadata_Treatment', 'Metadata_Plate_Well', 'Metadata_Plate','Metadata_Well']

# Create a new DataFrame with the selected columns
final_wells = most_common_rows.loc[:, selected_columns].copy()

final_wells.head(20)
final_wells.to_csv("final_wells.csv", index=False)


# In[114]:


#Choose treatments of interest to keep for site-level representative image identification
treatments_to_keep = ["F--IPL-A","F----LS-","F--I--S-","F-EI-L-A","F---PLS-","F--I----","F--IPLS-","F-E-----","FTEIPLSA","-TEI-L-A","GM-MSC","--------"]

final_wells_filt = final_wells[final_wells['Metadata_Treatment'].isin(treatments_to_keep)]
final_wells_filt.head(5)
final_wells_filt.to_csv("final_wells_filtered.csv", index=False)


# ## Step 2 - Use representative wells to find most representative Site images
# 
# In this step, Site-median values are calculated for each representative well and each feature of interest. The average of site-median values is calculated and the site-median that lies closest to the average is isolated and used as aggregate representative image

# In[115]:


# Filter single-cell dataframe to only include representative wells found in previous step
full_data_filt = full_data.merge(final_wells_filt[['Metadata_Plate_Well']], on='Metadata_Plate_Well', how='inner')
print(full_data_filt.shape)
full_data_filt.head(10)
#full_data_filt.to_csv("full_data_filt.csv", index=False)


# In[116]:


# Perform the aggregation - output site level median profiles
site_medians = aggregate(
    full_data_filt,
    strata=["Metadata_Plate", "Metadata_Well","Metadata_Site"],
    features="infer",
    operation="median",
    subset_data_df="none",
)
print("Site_Median profile size (row,col):", site_medians.shape)
site_medians.head(12)

# Make Plate_Well column: 
site_medians['Metadata_Plate_Well'] = site_medians['Metadata_Plate'].astype(str) + site_medians['Metadata_Well']
#medians = pd.merge(medians,N,on='Plate_Well',suffixes=('','_remove'))

# Remove duplicate columns: 
site_medians.drop([i for i in site_medians.columns if 'remove' in i], axis=1, inplace=True)
print("Shape of medians and cell count after removing duplicate cols: ", site_medians.shape)
#cols = ['size','Plate_Well','Metadata_Plate','Metadata_Well']
#medians = medians[cols + [c for c in medians.columns if c not in cols]]
site_medians.head(12)


# In[117]:


# Initialize an empty dictionary to store the results
isolated_sites = {}

for column in measurement_columns:
    # Calculate the mean of sites for each feature and unique Plate_Well
    mean_values = site_medians.groupby('Metadata_Plate_Well')[column].mean().reset_index()
    
    # Merge the mean values back into the original DataFrame
    site_medians = pd.merge(site_medians, mean_values, on='Metadata_Plate_Well', suffixes=('', '_mean'))
    
    # For each row, Calculate the absolute difference between the measurement and the mean of each treatment group
    site_medians[column + '_diff'] = abs(site_medians[column] - site_medians[column + '_mean'])
    
    # Find the 'Plate_Well' with the minimum absolute difference for each 'Metadata_Treatment' group
    isolated_sites[column] = site_medians.loc[site_medians.groupby('Metadata_Plate_Well')[column + '_diff'].idxmin()]

print("Calculations added to site_medians.csv size (row,col): ", site_medians.shape)
site_medians.head(16)
#site_medians.to_csv("site_medians.csv", index=False)


# In[118]:


# Create a DataFrame that combines all closest 'Plate_Well' results
isolatedSites_df = pd.concat(isolated_sites, axis=0, ignore_index=True)

print("Shape of dataframe with only minimum absolute difference well remaining (row,col): ", isolatedSites_df.shape)
isolatedSites_df.head(300)


# In[119]:


# Count the occurrences of each 'Site' for each 'Plate_Well'
count = isolatedSites_df['Metadata_Plate_Well'].value_counts().reset_index()
count.columns = ['Metadata_Plate_Well', 'Count']

# Merge the "count" DataFrame with the "isolatedSites_df" DataFrame
isolatedSites_df = isolatedSites_df.merge(count, on='Metadata_Plate_Well')

# Group the "isolatedSites_df" DataFrame by 'Metadata_Plate_Well' and find the row with the highest "Count" for each group
most_common_rows = isolatedSites_df.loc[isolatedSites_df.groupby('Metadata_Plate_Well')['Count'].idxmax()]

# List of column names you want to keep in "isolatedSites_df"
selected_columns = ['Metadata_Plate_Well', 'Metadata_Plate','Metadata_Well','Metadata_Site']

# Create a new DataFrame with the selected columns
final_sites = most_common_rows.loc[:, selected_columns].copy()

final_sites.head(20)
final_sites.to_csv("final_sites.csv", index=False)
```
